# Identity-by-descent segments in large samples

**DOI:** 10.1101/2024.06.05.597656

**Authors:** Seth D. Temple, Elizabeth A. Thompson

## Abstract

If two haplotypes share the same alleles for an extended gene tract, these haplotypes are likely to be derived identical-by-descent from a recent common ancestor. Identity-by-descent segment lengths are correlated via unobserved ancestral tree and recombination processes, which commonly presents challenges to the derivation of theoretical results in population genetics. We show that the proportion of detectable identity-by-descent segments around a locus is normally distributed when the sample size and the scaled population size are large. We generalize this central limit theorem to cover flexible demographic scenarios, multi-way identity-by-descent segments, and multivariate identity-by-descent rates. We use efficient simulations to study the distributional behavior of the detectable identity-by-descent rate. One consequence of non-normality in finite samples is that a genome-wide scan looking for excess identity-by-descent rates may be subject to anti-conservative control of family-wise error rates.

**Highlights:**

- We show the asymptotic normality of the detectable identity-by-descent rate, a mean of correlated binary random variables that arises in population genetics studies.
- We generalize our main central limit theorem to cover scenarios of nonconstant population sizes, multi-way identity-by-descent segments, and identity-by-descent rates of multiple samples from the same population.
- In enormous simulation studies, we use an efficient algorithm to characterize distributional properties of the detectable identity-by-descent rate.

## 1. Introduction

Two individuals share a haplotype segment identical-by-descent (IBD) if they inherit it from the same common ancestor. Here, we study the length of IBD segments that overlap a single focal location. Ignoring gene conversion, IBD segments are randomly cut by crossover recombination in each future generation. The length of an IBD segment is thus shorter with higher probability the more removed its common ancestor is from the present day.

Using modern methods, long IBD segments can be detected with high accuracy from genetic data [21, 39, 44, 57]. Detectable segments can provide rich information about the recent genetic history of a population sample. For instance, detected IBD segments have been used to test for rare variant associations when a disease allele is untyped or a genome-wide association study is underpowered [7, 23, 35]. They have also been used to estimate relatedness [21, 39, 57], haplotype phase [2, 33], mutation rates [37, 51, 52], recombination rates [55], gene conversion rates [6, 37], demographic changes [4, 8, 36], and positive selection [50]. We will study a sample mean of indicators if an IBD segment is long enough to be reliably detected. The binary random variables are correlated via unobserved recombinations and a random ancestral tree.

For independent, identically distributed data, maximum likelihood estimators are asymptotically consistent, efficient, and normally distributed under regularity conditions [10]. Composite likelihood approaches are commonly used in genetics when it is analytically intractable or computationally expensive to address dependencies in the data [30]. To what extent consistency, efficiency, and asymptotic normality extend to maximum composite likelihood estimators is generally unknown [30]. Studying maximum composite likelihood estimators can be especially challenging if their maxima do not have a closed form [36, 51]. In our work, the composite likelihood will be the binomial likelihood, which is maximized by the sample mean of binary random variables. The statistical property we care the most about is asymptotic normality, which is that the estimator’s distribution converges to a Gaussian distribution as the sample size tends to infinity [10].

Without theoretical results, some authors assume that their estimators are distributed within some parametric family. In one example, Palamara et al. [38] assume without proof that their estimator of coalescent rates within the past tens of generations is Gamma distributed. In another example, Carmi et al. [9] observe that the Gaussian distribution is a good fit for the average fraction of the genome shared IBD by an individual with any other individual. Still, this observation is not the same as a theoretical result. When the sampling distribution is not sub-normal [53], statistical inference assuming normality may understate the probability of extreme values.

Creating valid confidence intervals can be more straightforward when an estimator is asymptotically normally distributed. The parametric bootstrap approach proposed in Temple et al. [50] gives adequate coverage in selection coefficient estimation for numerous simulation studies. Their technique implicitly assumes that the rate of detectable IBD segments around a locus, and certain functions^1^ thereof, are normally distributed in large samples. In contrast, bootstrap resampling [16] has been employed in IBD-based estimation procedures [4, 6, 8, 36, 51]. For significance level *α*, these existing works do not demonstrate that their (1 − *α*)% bootstrap confidence intervals contain a true parameter in (1 − *α*)% of simulations. Moreover, nonparametric bootstrapping tends to give confidence intervals that are not wide enough to satisfy coverage [34].

Here, we derive sufficient conditions under which the proportion of detectable IBD segments around a locus is asymptotically normally distributed. The proof is to show that the variance of detectable IBD segments dominates the covariance between detectable IBD segments. Our conditions involve a minimum length of detectable IBD segments times the population size from which a large sample is drawn. The large population size requirement, in particular, indicates that most of the branch lengths in the ancestral tree must be long for the result to hold. The overall contribution of this work is to support IBD-based statistical inference with rigorous theory and extensive simulation studies.

The outline of the paper is as follows. In Section 2, we formally define our probability model for IBD segments that overlap a fixed location. In Section 3, we present and prove our main result for the asymptotic normality of the detectable IBD rate around a fixed location. In Section 4, we generalize our central limit theorem to cover nonconstant population sizes, multi-way IBD segments, and IBD rates between samples from the same population. In Section 5, we use simulation to investigate the statistical properties of IBD-based estimators and IBD graphs around a locus. Many calculations of covariance terms are left to the Appendix.

## 2. Preliminary material

First, we define our mathematical notation. The notation in Sections 2.1 and 2.2 follow the notation used in Temple et al. [49]. We use the Kingman coalescent [26, 27] as a model for the times until recent common ancestors, and we use the Poisson process to model recombination without interference. The probability that an IBD segment is longer than a detection threshold is derived by integrating over these two waiting time distributions.

### 2.1 The time until a common ancestor

Let *n* be the haploid sample size and *k* ≤ *n* be the size of a subsample. Define *N* to be the constant population size and *N*(*t*) the population size *t* generations ago. Let the random variable *T*_*k*_ denote the time until a common ancestor is reached for any two of *k* haploids, which we measure in units of *N* generations. In the discrete-time Wright-Fisher (WF) process, each haploid has a haploid ancestor in the previous generation, and if haploids have the same haploid ancestor, their lineages join.

The Kingman coalescent comes from the continuous-time limit of the WF process when subsample sizes are much smaller than constant population sizes. Specifically, *T*_*k*_ converges weakly to Exponential 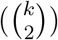 for *k* ≪ *N* and *N* →∞ [26, 27], where 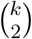 is the rate parameter. We focus on the times *T*_4_ ∼ Exponential(6), *T*_3_ ∼ Exponential(3), and *T*_2_ ∼ Exponential(1) until any two of four, three, and two haploids reach a common ancestor, respectively.

### 2.2 The distance until crossover recombination

The genetic distance (in Morgans) between two loci is the number of crossovers expected to occur in an offspring gamete. Assuming no interference in double-stranded breaks and independent crossovers, Haldane [24] derives that the genetic distance until a crossover recombination is exponentially distributed. This result leads to modeling crossover points along the genome as a Poisson process. Browning [3] considers crossover models without and with interference [28] when studying transitions between IBD states, whereas we exclusively use the model without interference.

From a fixed point, the Morgans distance in one direction until a gamete off-spring crossover is exponentially distributed with rate parameter 1. After *t* in-dependent meioses, the surviving haplotype segment length to the right of the focal location is distributed as Exponential(*t*), where *t* is the rate parameter. Let *a* and *b* be sample haplotypes in the current generation, and define *L*_*a*_, *R*_*a*_ | *t* ∼ Exponential(*t*) to be sample haplotype *a*’s recombination endpoints to the left and right of a focal location after *t* generations. Because crossovers to the left and right of the focal location are independent, the extant width from the ancestor at time *t* is *W*_*a*_ := *L*_*a*_ + *R*_*a*_ | *t* ∼ Gamma(2, *t*). Since the *t* meioses descend independently to *a* and *b* from their most recent common ancestor, the IBD segments that are shared by *a* and *b* are *L*_*a,b*_, *R*_*a,b*_ |*t* ∼ Exponential(2*t*) and *W*_*a,b*_ | *t* ∼ Gamma(2, 2*t*).

### 2.3 The presence of detectable IBD segments

Relative to a focal point, we consider the detection of long IBD segments in a sample. Let *X*_*a,b*_ := *X*_*a,b*_(*w*) = *I*(*R*_*a,b*_ ≥ *w*) indicate if the IBD segment to the right that is shared by sample haplotypes *a* and *b* is longer than a detection threshold *w* Morgans. The binary random variables {*X*_*a,b*_} are identically distributed with the same mean 𝔼_2_[*X*_*a,b*_] and correlated through the unobserved coalescent tree. We use 𝔼_2_, 𝔼_3_, and 𝔼_4_ and Cov_2_, Cov_3_, and Cov_4_ to denote expected values and covariances with respect to coalescent trees of two, three, and four sample haplotypes, respectively.

Our central limit theorem concerns a mean of the IBD segment indicator random variables. Namely, the detectable IBD rate to the right of a fixed location is

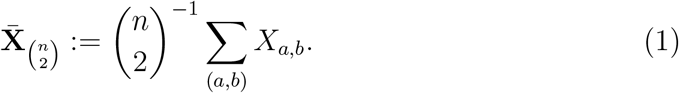

Let *Z*_*a,b*_ := *X*_*a,b*_ − 𝔼_2_[*X*_*a,b*_] be the mean-centered binary random variable, and let the sum of all except one of these mean-centered random variables be **Z**_−*a,b*_ := Σ_(*c,d*)_ *Z*_*c,d*_ − *Z*_*a,b*_. The sum of variances of all IBD segment indicators is

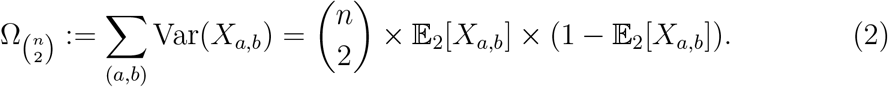

Finally, the mean-centered and suitably scaled detectable IBD rate to the right of a locus is

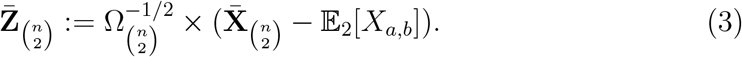

For IBD segments overlapping a focal location, let *Y*_*a,b*_ := *I*(*L*_*a,b*_ + *R*_*a,b*_ ≥ *w*) and 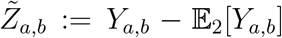. The terms 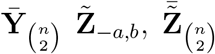, and 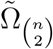, are defined analogously to 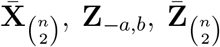, and 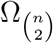, respectively. We drop the subscript 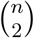 when it is clear that the aggregation is over 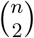 pairs of haplotypes. Figure 1 provides a conceptual example calculating 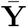 for four sample haplotypes.

**Figure 1.**
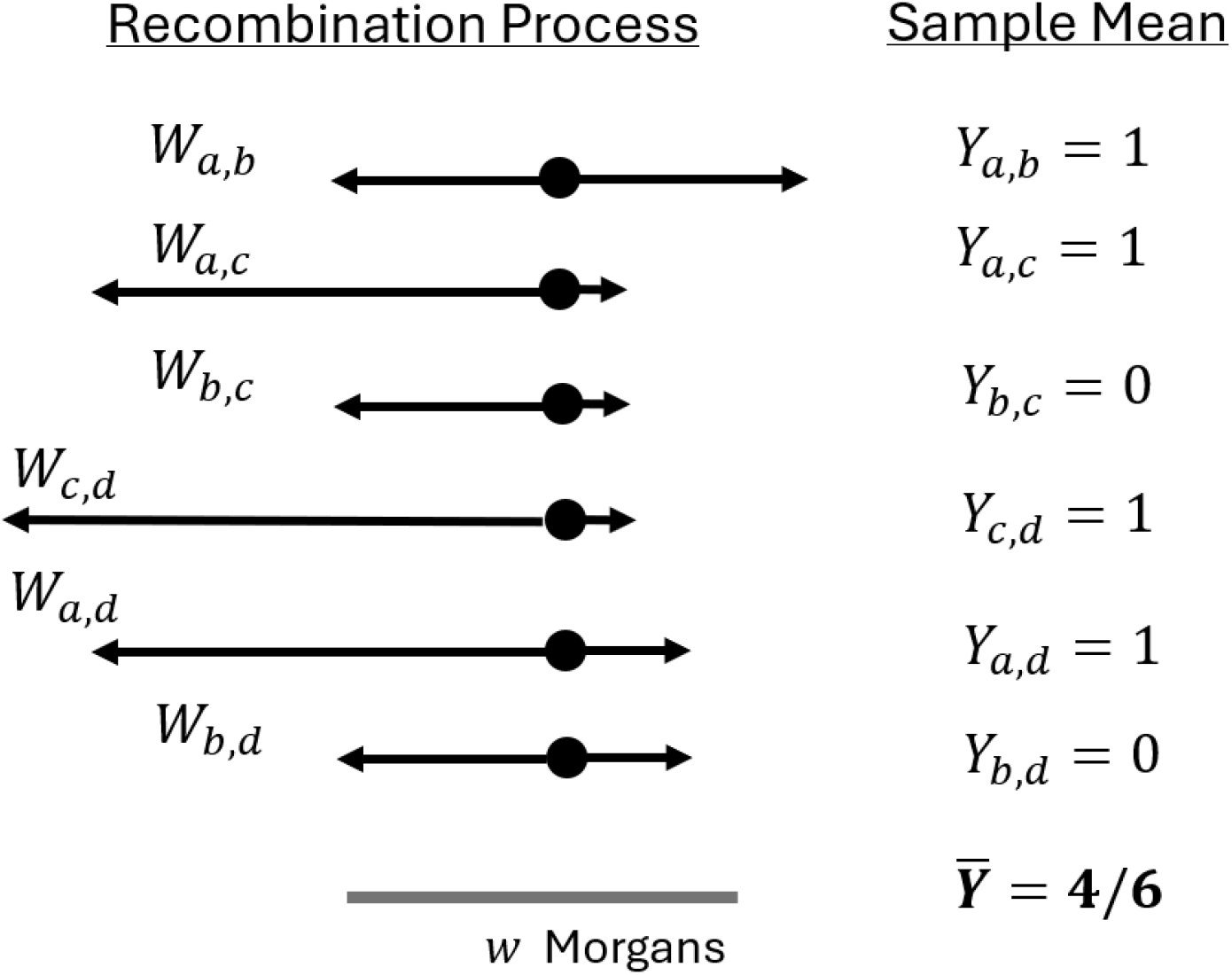
Example calculation of the detectable IBD rate. IBD segment lengths overlapping a focal point for sample haplotypes *a, b, c, d* are shown. The IBD segment indicators (*Y*_*i,j*_’s) are 1 if their IBD segment lengths (*W*_*i,j*_’s) exceed *w* Morgans and otherwise 0. The detectable IBD rate 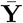 is the mean of these correlated binary random variables. The detectable IBD rate to the right of the focal point, 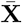, is calculated similarly.

We use additional subscript indices when segments are IBD among multiple haplotypes, which we refer to as multi-way IBD segments. For instance, *Y*_*a,b,c*_ indicates if the IBD segment around a locus that is shared between haplotypes *a, b*, and *c* is longer than *w* Morgans. The corresponding sample mean over 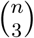 haplotype triplets is denoted 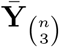, and the related sums, means, and variances are defined similarly. This notation is important to extend our main central limit theorem to multi-way IBD segment indicators.

We use the superscript *l* to denote the sample label when different population samples are considered. For example, 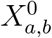 and 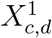 indicate if the IBD segments around a locus that are shared between haplotypes *a* and *b* in population sample 0 and *c* and *d* in population sample 1 are longer than *w* Morgans, respectively. Mean-centered and bold-faced terms are defined analogously for these extensions. For example, the mean in population sample 0 of 2-way IBD segment indicators overlapping a focal location is denoted 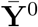. This notation is important to extend our main univariate central limit theorem to a multivariate Gaussian version.

## 3. Main central limit theorem

If *U*_1_, …, *U*_*n*_ ∼^*iid*^ *G* for some model *G*, the Lindeberg-Lévy central limit theorem says that the standardized sample mean weakly converges to the standard normal distribution (under some regularity conditions) [31]. The special case of this result for binary random variables [15] is more closely related to our work. The result does not apply in our case because the IBD segment indicators {*X*_*a,b*_} to the right of a focal point are not independent. We start by focusing on the mean-centered and suitably scaled detectable IBD rate 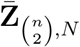 to the right of a focal location, where the subscript *N* clarifies that the haplotypes are sampled from a population of constant size *N*.

Our central limit theorems concern large sample size *n* and large population size *N* scaled by the Morgans detection threshold *w*. The intuition for our weak law is that the covariance between IBD segment indicators Σ _(*a,b*)≠(*c,d*)_ Cov(*X*_*a,b*_, *X*_*c,d*_) is small relative to the sum of the variances of the individual IBD segment indicators 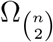. The sum of covariances between random variables being negligible compared to the sum of variances of the random variables themselves is the basis of the general central limit theorem for dependent data that is given in Chandrasekhar and Jackson [11] and Chandrasekhar et al. [12].

### Theorem 3.1

*For n and Nw tending to infinity, the mean-centered and suitably scaled detectable IBD rate* 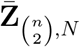, *to the right of a focal location converges in distribution to the standard normal distribution when the following are true:*

1. *Nw* = *o*(*n*^2^), *scaled population size is small relative to the number of pairs;*
2. *n* = *o*(*Nw*), *sample size is small relative to scaled population size;*
3. 𝔼[*Z*_*a,b*_ *×* **Z**_−*a,b*_|**Z**_−*a,b*_] ≥ 0 *for all* **Z**_−*a,b*_.

*Proof*. We show that our three conditions are sufficient to apply Corollary 1 in Chandrasekhar et al. [12]. Without loss of generality, we derive integrals over a tree with two sample haplotypes *a* and *b*, a tree with three sample haplotypes *a, b*, and *c*, and a tree with four sample haplotypes *a, b, c*, and *d*.

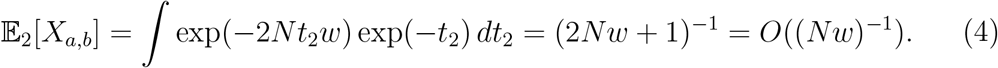

It is easy to show that 𝔼_2_[*X*_*a,b*_] *→* 0 uniformly for large scaled population size (Lemma A.1). The second condition implies that 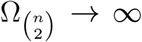. The assumption in Chandrasekhar et al. [12] that 𝔼[|*Z*_*a,b*_|^3^]*/*𝔼[|*Z*_,*ab*_|^2^]^3*/*2^ is bounded above is true for nondegenerate Bernoulli random variables [11] (Lemma A.2). Lastly, given *n* = *o*(*Nw*), we show that

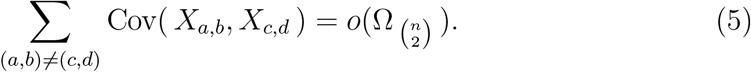

In Appendix A.1, we derive bounds on the integrals Cov_3_(*X*_*a,b*_, *X*_*a,c*_) = *O*((*Nw*)^−2^) and Cov_4_(*X*_*a,b*_, *X*_*c,d*_) = *O*((*Nw*) ^−3^). Next, there are *n*(*n* − 1)(*n* − 2) ∼ *n*^3^ combinations of three haplotypes *a, b*, and *c*, and there are *n*(*n* − 1)(*n* − 2)(*n* − 3)/4 ∼ *n*^4^ combinations of four haplotypes *a, b, c*, and *d*. In asymptotic arguments, the notation ∼ means asymptotic equivalence, not distributed as.

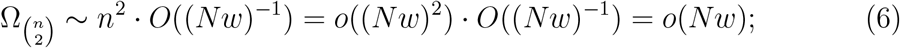

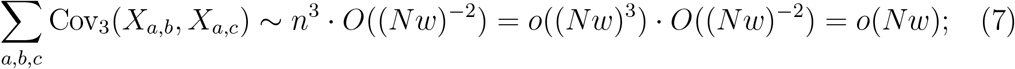

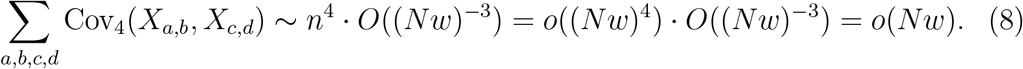

The covariance between IBD segment indicators (Equations 7 and 8) is controlled by the covariance within IBD segment indicators (Equation 6).

The first two conditions have appealing interpretations. First, *Nw* = *o*(*n*^2^) says that the sample size is large enough relative to the scaled population size such that we observe many IBD segments to the right of a focal location that are longer than the Morgans threshold *w*. Second, *n* = *o*(*Nw*) says that the sample size is not too large relative to the scaled population size such that we do not observe many large clusters of haplotypes with IBD segments to the right of a focal location that are longer than the Morgans threshold *w*.

The third condition also has an interpretation in the context of population genetics. It says that if the number of detectable IBD segments to the right of a focal location, except for *X*_*a,b*_, is less than the expectation 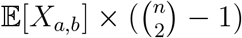, then the IBD segment to the right of a focal location that is shared by *a* and *b* is shorter than *w* Morgans on average, and vice versa if **X**_−*a,b*_ is greater than its expected value. This assumption seems plausible if IBD segments to the right of a focal location have nonnegative covariance, which we show in Appendix A.1. Moreover, one intuits that the posterior distribution of *X*_*a,b*_|**X**_−*a,b*_ is more likely to come from a tree with long branches than the unconditional distribution of *X*_*a,b*_ is when 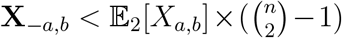, and vice versa when 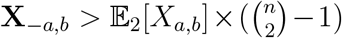.

One can show that the small sample size *n* = 3 is a pathological example where the third condition breaks down (Lemma A.6). We do not otherwise calculate 𝔼[*Z*_*a,b*_ *×* **Z**_−*a,b*_|**Z**_−*a,b*_] for all **Z**_−*a,b*_, which involves integration over the space of all coalescent trees and the 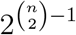 hypercube of 0’s and 1’s. In a simulation study, we evaluate the third condition via the Monte Carlo method (Appendix A.2), concluding that this condition likely holds in large samples.

The asymptotic normality of 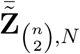 follows from the same arguments as those of the proof in Theorem 3.1. We show in Appendix A.1 that Cov_2_(*Y*_*a,b*_, *Y*_*a,b*_), Cov_3_(*Y*_*a,b*_, *Y*_*a,c*_), and Cov_4_ (*Y*_*a,b*_, *Y*_*c,d*_) are *O*((*Nw*)^−1^), *O*((*Nw*)^−2^), and *O*((*Nw*) ^−3^), respectively.

### Theorem 3.2.

*For n and Nw tending to infinity, the mean-centered and suitably scaled detectable IBD rate* 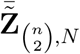 *around a locus converges in distribution to the standard normal distribution when the following are true:*

1. *Nw* = *o*(*n*^2^);
2. *n* = *o*(*Nw*);
3. 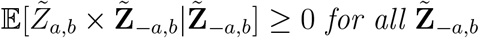.

## 4. Extensions

### 4.1 Flexible demographic scenarios

We can derive a similar result for varying population sizes. Let *N*_1_ = max_*t*_ *N*(*t*) and *N*_2_ = min_*t*_ *N*(*t*). Compared to varying population sizes *N*(*t*), the indicator of a detectable IBD segment around a focal location has larger expected value and variance when sample haplotypes come from a constant population of size *N*_2_. Conversely, compared to varying population sizes *N*(*t*), the indicator of a detectable IBD segment around a focal location has smaller expected value and variance when sample haplotypes come from a constant population of size *N*_1_. We use these facts to establish covariance bounds for complex demography.

#### Theorem 4.1

*For n, N*_1_*w, and N*_2_*w tending to infinity, rhe mean-centered and suitably scaled detectable IBD rate* 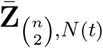 *to the right of a focal location converges in distribution to the standard normal distribution when the following are true:*

- *N*_1_*w* = *o*(*n*^2^);
- *n* = *o*(*N*_2_*w*);
- 𝔼[*Z*_*a,b*_ *×* **Z**_−*a,b*_|**Z**_−*a,b*_] ≥ 0 *for all* **Z**_−*a,b*_.

*The same conditions imply weak convergence for* 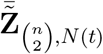

*Proof*. The argument is the same as in Theorem 3.1, except we use *N*_1_ and *N*_2_ to upper and lower bound covariance terms.

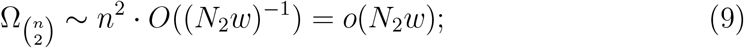

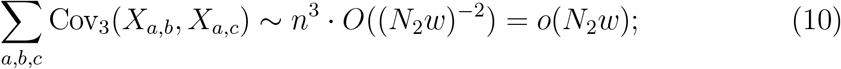

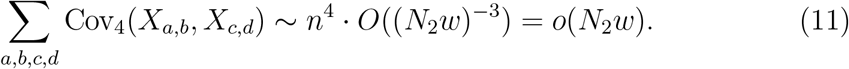

Theorem 3.1 is a special case of Theorem 4.1 when *N*_1_ = *N*_2_. The conditions in Theorem 4.1 are unlikely to hold in real data examples and are more difficult to interpret. Note that the proof of Theorem 4.1 does not make use of the entire curve *N*(*t*). The population sizes at the most recent coalescent times impact the covariance of and between IBD segments around a focal location the most. As in Theorem 3.2, we can extend Theorem 4.1 to address IBD segments overlapping a focal location.

### 4.2 Multi-way IBD segments

To calculate the probability that an *m*-way IBD segment indicator is 1, we integrate over *m* − 1 coalescent times and the recombination processes at these common ancestors. Here, we consider *m >* 2 but *m* much smaller than the sample size *n*. For example, we compute the expected value of the 3-way IBD segment indicator to the right of a focal location

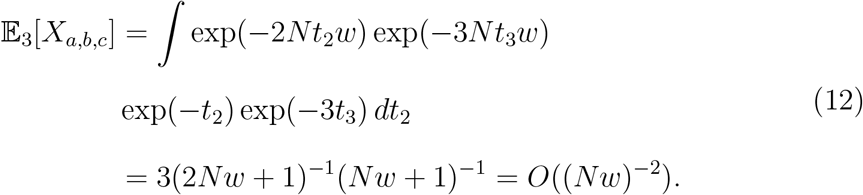

Note in this derivation and that of Equation 4 fall under the general result that 𝔼_*m*_[*X*_…*m*_] = *O*((*Nw*)^−(*m*−1)^), where … *m* denotes *m* labeled haplotypes. To observe many *m*-way IBD segment indicators, we require (*Nw*)^*m*−1^ = *o*(*n*^*m*^) because the sums are over 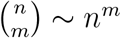 identically distributed random variables.

#### Theorem 4.2

*For n and Nw tending to infinity and bounded m* = *O*(1), *the mean-centered and suitably scaled detectable IBD rate* 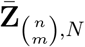 *to the right of a focal location converges in distribution to the standard normal distribution when the following are true:*

- (*Nw*)^*m*−1^ = *o*(*n*^*m*^);
- *n* = *o*(*Nw*);
- 𝔼[*Z*_…*m*_ *×* **Z**_−…*m*_|**Z**_−…*m*_] ≥ 0 *for all* **Z**_−…*m*_.

*The weak convergence result holds for* 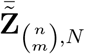 *under the same conditions*.

*Proof*. The proof is again to show that the three conditions are sufficient to apply Corollary 1 in Chandrasekhar et al. [12]. The strategy is to calculate the relevant integrals 𝔼_*m*_[·], …, 𝔼_2*m*_[·], count the number of occurrences of each covariance type, and then observe that the condition *n* = *o*(*Nw*) is sufficient to control the total covariance. In Appendix A.1.2, we give a full proof for the 3-way IBD rate, from whose covariances and combinatorics it is straightforward to see a pattern as *m* increases.

Theorems 3.1 and 3.2 are special cases of Theorem 4.2 when *m* = 2. We remark that *n* = *o*(*Nw*), which does not involve *m*, is a condition shared between Theorems 3.1 and 4.2. Recall that this condition maintains that covariances between IBD segment indicators are small, which is governed by large scaled population size *Nw*.

### 4.3 Multivariate IBD rates

We now show that the conditions *n* = *o*(*Nw*) and *Nw* = *o*(*n*^2^) are also sufficient to apply the multivariate version of the Chandrasekhar et al. [12] central limit theorem. From the multivariate result, we can derive the asymptotic distribution of the difference in IBD rates between case and control sample sets. To extend our main result to multivariate random vectors, we consider the example of two disjoint sample sets labeled 0 and 1. Each sample consists of *n* samples from the same population of size *N*.

Let 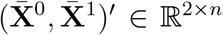 be the column vector of two sample means, where ^*′*^ is transpose. The detectable identity-by-descent segment rates around a locus are denoted 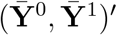, and the standardized sample means are denoted 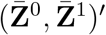 and 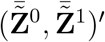. In general, we denote 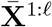 and 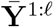 and 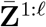 and 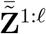 as the unstandardized and standardized IBD rates to the right of and overlapping a focal location for *l* distinct samples of *n* haplotypes from *N*. The mean-centered sums of IBD segment indicators excluding 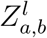 and 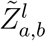 are denoted 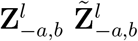, respectively.

#### Theorem 4.3

*For bounded* 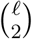 *and n and Nw tending to infinity, the mean-centered and suitably scaled IBD rates* 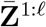 *converge in distribution to the standard normal distribution N*_*ℓ*_ (**0, I**_*ℓ*×*ℓ*_) *when the following are true:*

- *Nw = o(n2);*
- *n = o(Nw);*
- 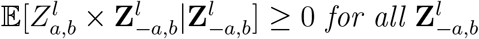.

*The weak convergence result holds for* 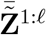 *under the same conditions*.

*(The proof is in Appendix A.1.3 using the result from Chandrasekhar et al. [12].)*

One important consequence of Theorem 4.3 is that affine transformations of the sample means column vector are asymptotically normally distributed. In particular, for the example of two samples and the row vector (1, −1), the difference in standardized IBD rates around a locus 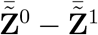 is asymptotically normally distributed. When there are *l* sample sets, for each pair of the *l* sample means, a row vector exists such that the dot product gives the difference in their IBD rates.

To apply Corollary 1 of Chandrasekhar et al. [12], we restrict our result to equally sized samples of *n* haplotypes. In case-control studies, there may be samples of unequal sizes *n*_1_ and *n*_0_. We conjecture that the difference in IBD rates will still asymptotically normally distributed, so long as 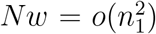 and 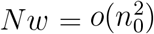 and max(*n*_0_, *n*_1_) = *o*(*Nw*). The conditions 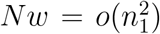 and 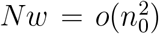 maintain that we detect many IBD segments in both samples. The condition max(*n*_0_, *n*_1_) = *o*(*Nw*) maintains that covariances are vanishing both in the diagonal terms 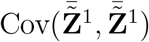 and 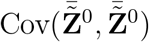 and the off-diagonal term 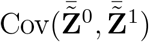.

Another limitation is our restriction to distinct sample sets, which is necessary to make the covariance calculations analytically tractable. Browning and Thompson [7] study the IBD rates between case-case, case-control, and control-control haplotype pairs, resulting in a sample means vector that does not fall under our mathematical framework. We conjecture that the empirical distributions of such vectors may be similar to those of vectors of nonoverlapping sample sets when the samples come from a large population. The reason for our conjecture is the same as before: the large scaled population size leads to vanishing covariances in the diagonal and off-diagonal terms.

## 5. Simulation studies

The theoretical results in Sections 3 and 4 rely on asymptotic conditions, not finite sample conditions. Using simulation, we explore the finite sample empirical distributions and percentiles of detectable IBD rate-based statistics around a fixed location. To investigate normality, we require massive simulations to form tens of thousands of empirical distributions.

We use the algorithm in Temple et al. [49] to simulate detectable IBD segments overlapping a fixed location. Despite the speed of the algorithm, the enormous scope of our simulations takes hundreds of days of computing time, which we spread across core processing units. If not for the algorithm’s efficiency, we would be limited in our ability to study the distributional behavior of the standardized detectable IBD rate 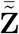 and the difference in IBD rates 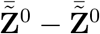.

We consider sample sizes of five and ten thousand “diploid” individuals. To implement “diploids”, we use a haploid model with two times the sample size of diploids (and likewise for demographic models). We consider the same demographic scenarios described in Temple et al. [50] and Temple et al. [49]: constant population sizes ranging from ten thousand to ten million diploid individuals and examples of exponential growth phases and a population bottleneck. Both complex demographic scenarios amount to population sizes ≥ 10^6^ in the most recent tens of generations and population sizes ≤ 10^4^ more than a few hundred generations ago. Figure S1 from Temple et al. [49] illustrates some of these demographic scenarios.

### 5.1. Identity-by-descent rates in finite samples

#### 5.1.1. Constant population sizes

Using the Shapiro-Wilk test [41, 42, 43], we investigate if empirical distributions of Σ_*a,b*_ *Y*_*a,b*_ resemble normal distributions as sample size *n*, population size *N*, and the Morgans length threshold *w* increase. We partition simulated IBD rates into five hundred empirical distributions based on one thousand observations. The null hypothesis is that the empirical distribution of detectable IBD rates is normally distributed. Rejecting the null hypothesis means that there is enough evidence indicating that the empirical distribution is not normal. We report the proportion of times we reject the null hypothesis at the significance level 0.05.

Figure 2 shows the proportion of rejected tests for increasing population size and Morgans length threshold with sample size fixed at five and ten thousand diploid individuals. The trend is that the proportion of rejected tests decreases with the increasing population size and Morgans length threshold. Figure S2 shows that this trend does not depend on the significance level. These observations align with the condition *n* = *o*(*Nw*) in Theorem 3.1 and Theorem 3.2. The setting for which the proportion is closest to 0.05 is *n* = 10^4^, *N* = 10^6^, and *w* = 0.04. Interestingly, for the same sample size and Morgans length threshold, we observe more rejected tests for *N* = 10^7^ than for *N* = 10^6^. This observation aligns with the condition *Nw* = *o*(*n*^2^) in Theorem 3.1 and Theorem 3.2 (there are too few observed IBD segments).

**Figure 2.**
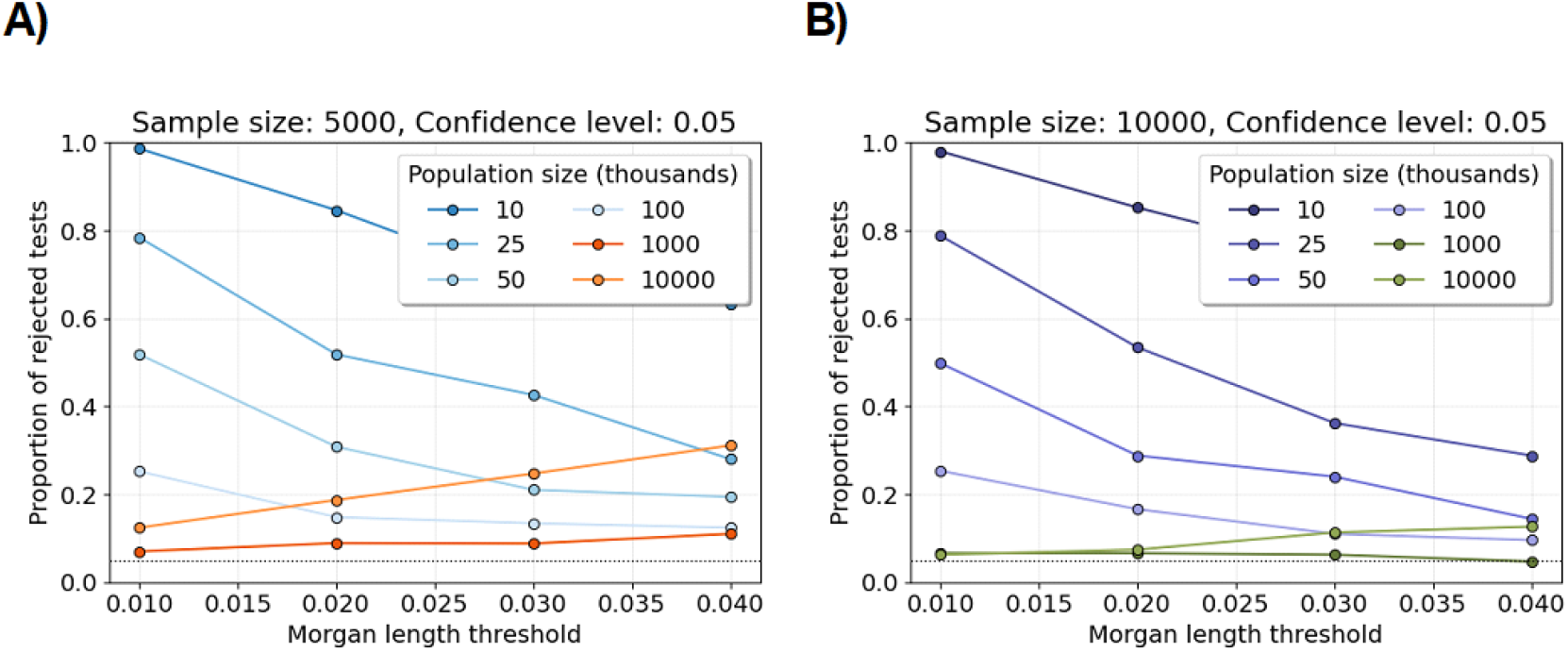
Shapiro-Wilk tests for varying population sizes. Line plots show the proportions of Shapiro-Wilk tests rejected at the significance level 0.05 (y-axis) for varying population size and fixed sample size. Each proportion is computed over five hundred tests. Each test is based on one thousand simulations of the number of identity-by-descent lengths longer than a specified Morgans length threshold (x-axis). A) The sample size is five thousand individuals. B) The sample size is ten thousand. The legends assign colors to different population sizes. The horizontal dotted line is at 0.05.

Figure S3 shows the proportion of rejected tests for increasing sample size and Morgans length threshold with population size fixed at fifty and one hundred thousand diploid individuals. The proportion of rejected tests decreases slightly with increasing sample size. This trend may be explained by the fact that sample size has no effect on the marginal correlations of IBD segments (Lemmas A.3, A.4, and A.5).

#### 5.1.2. Flexible demographic scenarios

Figure S4 shows the proportion of rejected tests for increasing sample size and Morgans length threshold in the three phases of exponential growth and population bottleneck demographic scenarios. For Morgans length threshold greater than or equal to 0.03, the proportions of rejected tests are less than 0.3 and 0.1 in the three phases of exponential growth and population bottleneck scenarios, respectively. Consistent with our central limit theorems, we observe a decreasing trend as we increase the Morgans length threshold, even though the proportions of rejected tests around 0.3 and 0.1 are not close to the nominal significance level 0.05. Additionally, these proportions are less than their corresponding proportions in the population of twenty-five thousand diploid individuals (Figure 2).

The conditions on the global extrema of population sizes in Theorem 4.1 are very stringent. The most recent population sizes have the strongest impact on the covariances of IBD segment indicators. One interpretation of the results in Figure S4 is that the detectable IBD rate around a locus may behave like a normal distribution in demographic scenarios with large recent population sizes, regardless of the not-so-recent population sizes.

#### 5.1.3. Difference of identity-by-descent rates in two samples

We compute the difference of detectable IBD rates around a locus by splitting five thousand diploid individuals into two equally sized subsets. Then, under different experimental conditions, we perform two hundred and fifty Shapiro-Wilk tests based on five hundred simulations of the test statistic.

Figure S4 shows the proportion of rejected tests for increasing population size and Morgans length threshold. At the significance level 0.05, and for all scaled population sizes, between 0.05 and 0.15 percent of tests are rejected. At the significance level 0.10, and for all scaled population sizes, between 0.10 and 0.30 percent of tests are rejected. There is no apparent trend as either population size or Morgans length threshold increases. One explanation is that any potential overdispersion of 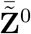 and 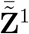, relative to the standard normal distribution, may be partially balanced out by considering the difference of the rates. Another explanation is the limited power to reject the Shapiro-Wilk null hypothesis in the scope of our computationally feasible experiments.

Across all simulation experiments in Sections 5.1.1, 5.1.2, and 5.1.3, we reject normality at rates greater than the Type 1 error rate 0.05 with the sample sizes and population sizes explored here. These magnitudes are already quite large relative to existing sample sizes and inferred effective population sizes. Nevertheless, the trends of increasing sample size and scaled population size suggest the validity of our central limit theorems.

### 5.2. Percentiles of the finite sample distributions

Next, we investigate possible explanations for rejecting the nominal significance levels at elevated rates. We focus on the upper percentiles of the empirical distribution of our test statistics 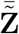 and 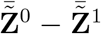. For each batch of simulations, we compute a mean, a standard deviation, and the mean plus three or four standard deviations. Then, we calculate the 99.86501^th^ and 99.99683^th^ percentiles of the test statistic over all batches. (These percentiles correspond to the standard normal quantiles of three and four.) We multiply the reciprocal of these quantiles by their corresponding estimated upper bounds, which we refer to as the relative upper bounds.

#### 5.2.1. The identity-by-descent rate in one sample

Browning and Browning [5], Temple et al. [50], and Temple [48] conduct hypothesis tests to evaluate if the detectable IBD rate 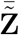 around any specific locus exceeds a genome-wide mean IBD rate. When our central limit theorems hold, we can interpret their hypothesis test as a one-sample one-sided z tests. Our estimated upper bounds, the mean plus some standard deviations, are meant to mimic their hypothesis tests [5, 50].

Figures 3 and S6 show the average relative upper bounds by increasing population size and Morgans length threshold. The average estimated upper bounds are less than the simulated percentile threshold for all sample sizes, population sizes, Morgans length thresholds, and quantiles considered. The average estimated upper bound is proportionally closer to the percentile threshold as population size and Morgans length threshold increase, which is a result consistent with Section 5.1.1 and our central limit theorems.

**Figure 3.**
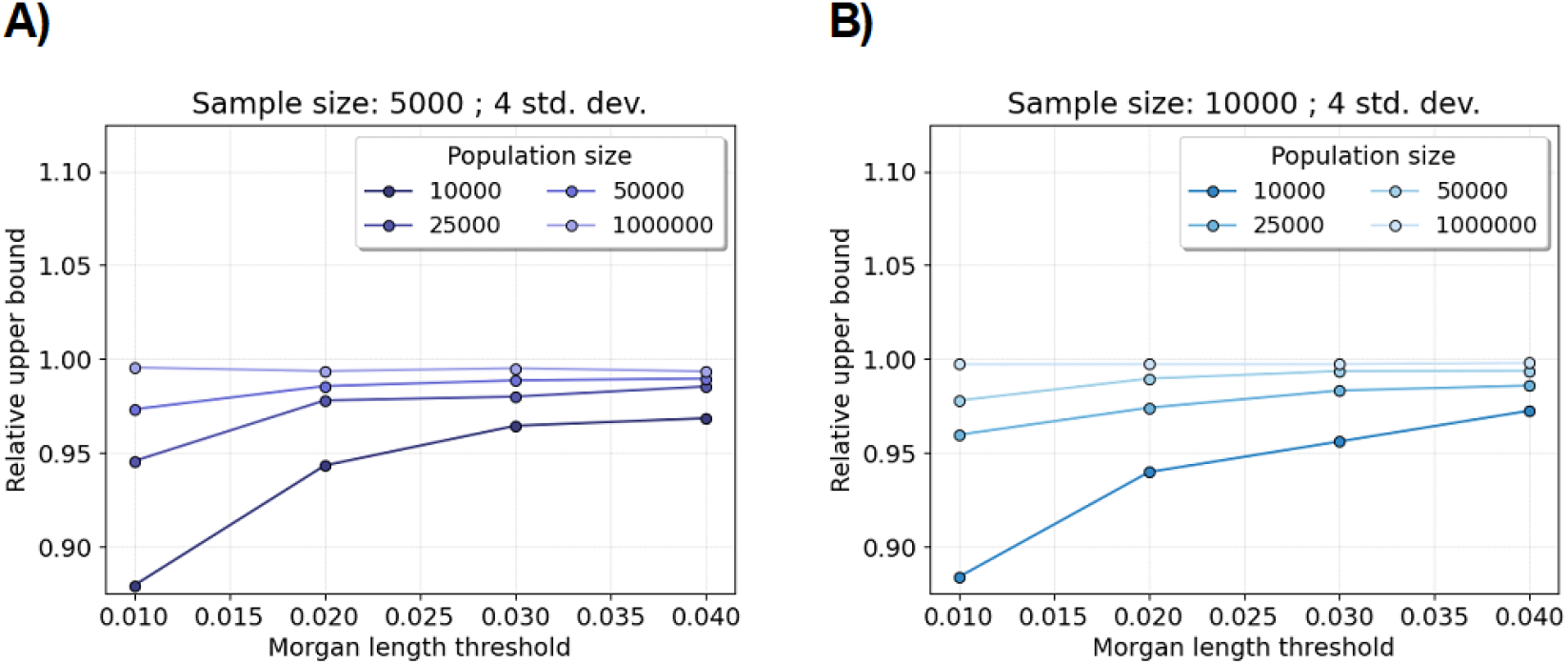
Relative upper bound for excess IBD scan. Line plots show the average mean plus four standard deviations divided by the 99.99683 percentile over two million simulations (y-axis). (The standard normal survival function of four is 0.9999683.) Each average relative upper bound is computed over one thousand tests. Each test is based on two thousand simulations of the number of identity-by-descent lengths longer than a specified Morgans length threshold (x-axis). A) The sample size is five thousand diploid individuals. B) The sample size is ten thousand diploid individuals. The legends assign colors to different constant population sizes.

Figure S4 shows that the average estimated upper bound is also less than the simulated percentile threshold for all sample sizes and Morgans length thresholds in the complex demographic scenarios. The average estimated upper bound is proportionally closer to the percentile threshold for the population bottleneck scenario compared to the three phases of exponential growth scenario, which is the complex demographic scenario with larger recent population sizes (Figure S1).

These experiments suggest that one reason why we reject the Shapiro-Wilk null hypothesis at elevated rates is because the test statistic’s upper tail probability is heavier than that of the standard normal distribution.

#### 5.2.2. Difference of identity-by-descent rates in two samples

Analogous to the excess IBD rate test, the difference in IBD rates 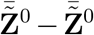 may be used as a hypothesis test for equality of means between two labeled subgroups. We perform the same experiment, except for the difference in IBD rates as our test statistic.

Figure S7 shows the average relative upper bounds by increasing population size and Morgans length threshold. We see no trend between the average relative upper bounds and sample size, population size, and Morgans length threshold, respectively. Compared to our observation in the one-sample experiment, the test statistic’s upper tail probability is not noticeably different from that of the standard normal distribution. These empirical observations are consistent with our Type 1 error experiment in Section 5.1.3.

### 5.3. Identity-by-descent graphs around a locus

Clusters of detectable IBD haplotypes overlapping a focal point indicate non-negligible covariance between segments. These cluster covariances could thus explain the observed non-normality in finite samples. We form detectable IBD graphs about a locus by drawing an edge between haplotypes if they share a detectable IBD segment overlapping a focal point. We define detectable IBD clusters as the connected components in the detectable IBD graph.

We analyze five features of graphs. The number of edges is equivalent to the number of IBD segments longer than the length threshold. A tree of order *m* is a connected component that has *m* nodes and *m* − 1 edges. An order *m* complete connected component has *m* nodes and edges between every pair of nodes. We count the number of trees of order 2 and 3, the number of complete connected components of order 3 or more, and the number of nodes in the largest connected component. We calculate the average, variance, minimum, and maximum for each feature over replicate simulations. We also conduct Shapiro-Wilk tests by splitting the simulated data as described in Section 5.1.1.

Note that the number of trees of order *m* is not the same as the *m*-way IBD rate around a locus. For example, in a complete connected component of four nodes, there are 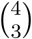 counts of 3-way detectable IBD. As a result, Theorem 4.2 does not apply to the following experiments on tree orders. However, we might expect to see some approximately normally distributed data if most components of degree *m* are trees.

#### 5.3.1. Comparing to sparse Erdős-Rényi graphs

The Erdős-Rényi graph is a simple network model in which independent edges between nodes occur with a uniform success probability [17]. We denote a sparse Erdős-Rényi network as one in which the success probability is vanishingly small. We compare the features of connected components between detectable IBD and Erdős-Rényi graphs, setting the uniform success probability to be the approximate probability of an IBD segment longer than a Morgans length threshold [36]. This contrast analyzes the evolution of independent edges versus weakly correlated edges of a specific nature.

For sparse Erdős-Rényi graphs, there are theoretical properties associated with the graph features that we consider in our simulation study. When the success probability is small, the number of trees of order m weakly converges to a Gaussian distribution in large networks [18], and trees of order m_1_ have faster convergence than trees of order m_2_ when m_1_ < m_2_. Another asymptotic property of sparse Erdős-Rényi graphs is that almost all nodes are in trees of small order or a single “giant” component [18].

Figure 4 shows that some empirical distributions of graph features resemble normal distributions in a sample size of five thousand diploid individuals from a population of one hundred thousand diploid individuals. Table 1 compares our summary statistics between these simulated detectable IBD and sparse Erdős-Rényi graphs. The variance, minimum, and maximum number of edges are larger for detectable IBD graphs compared to sparse Erdős-Rényi graphs, which is a direct consequence of the nonzero covariance of IBD edges^2^. The proportions of rejected hypothesis tests for numbers of trees of order 2 and connected components of degree 3 or more are close to 0.05 for both detectable IBD and sparse Erdős-Rényi graphs. While we observe that some limiting distributional behaviors of small degree connected components in detectable IBD graphs match those in sparse Erdős-Rényi graphs, these observations go beyond the theory we have presented.

**Table 1:**
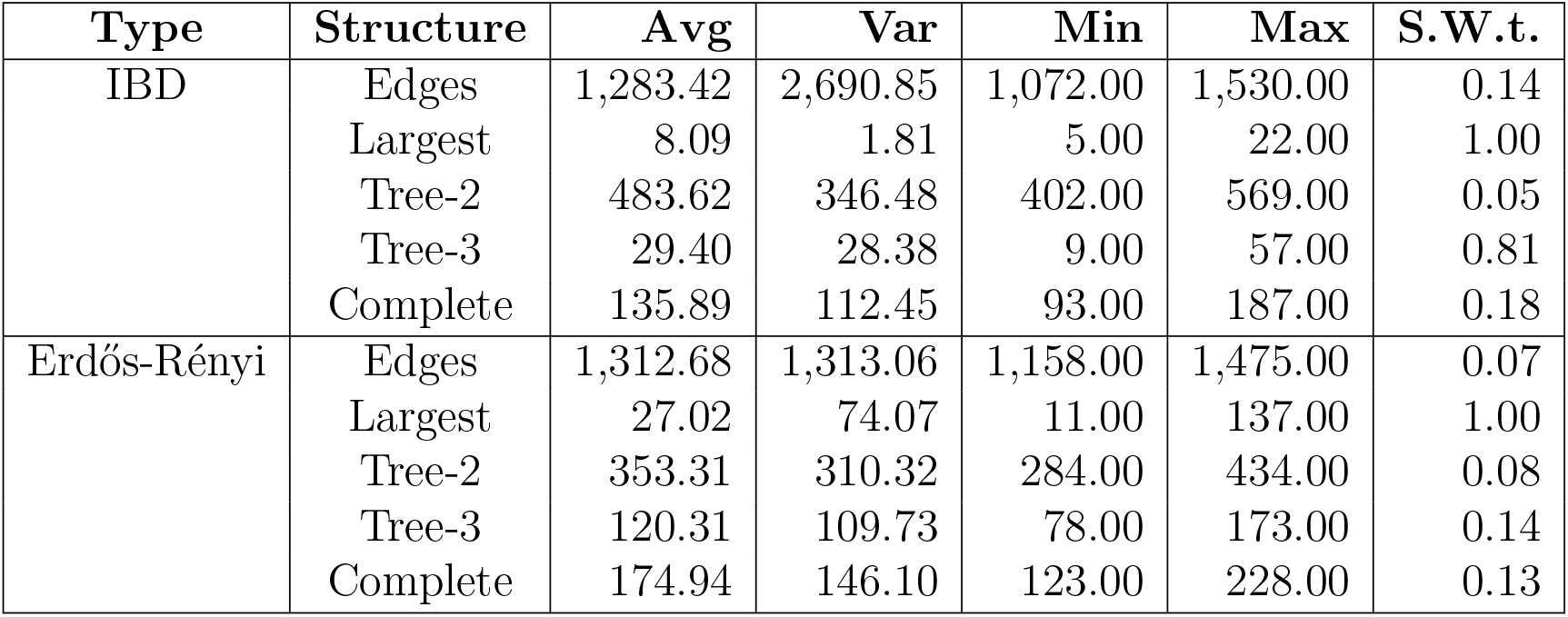
Summary statistics of IBD and Erdős-Rényi graphs. Network structures of interest are the number of edges (Edges), the degree of the largest components (Largest), the number of trees of order 2 and 3 (Tree-2 and Tree-3), and the number of complete components of degree 3 or more (Complete). Summary statistics are aggregated over 125,000 simulations. Shapiro-Wilk tests at the significance level 0.05 are performed with 500 replicates for 250 simulations, and the proportion of rejected null hypotheses reported as S.W.t. The population size is one hundred thousand diploid individuals. The sample size is two thousand diploid individuals. The Morgans length threshold is 0.03.

**Figure 4.**
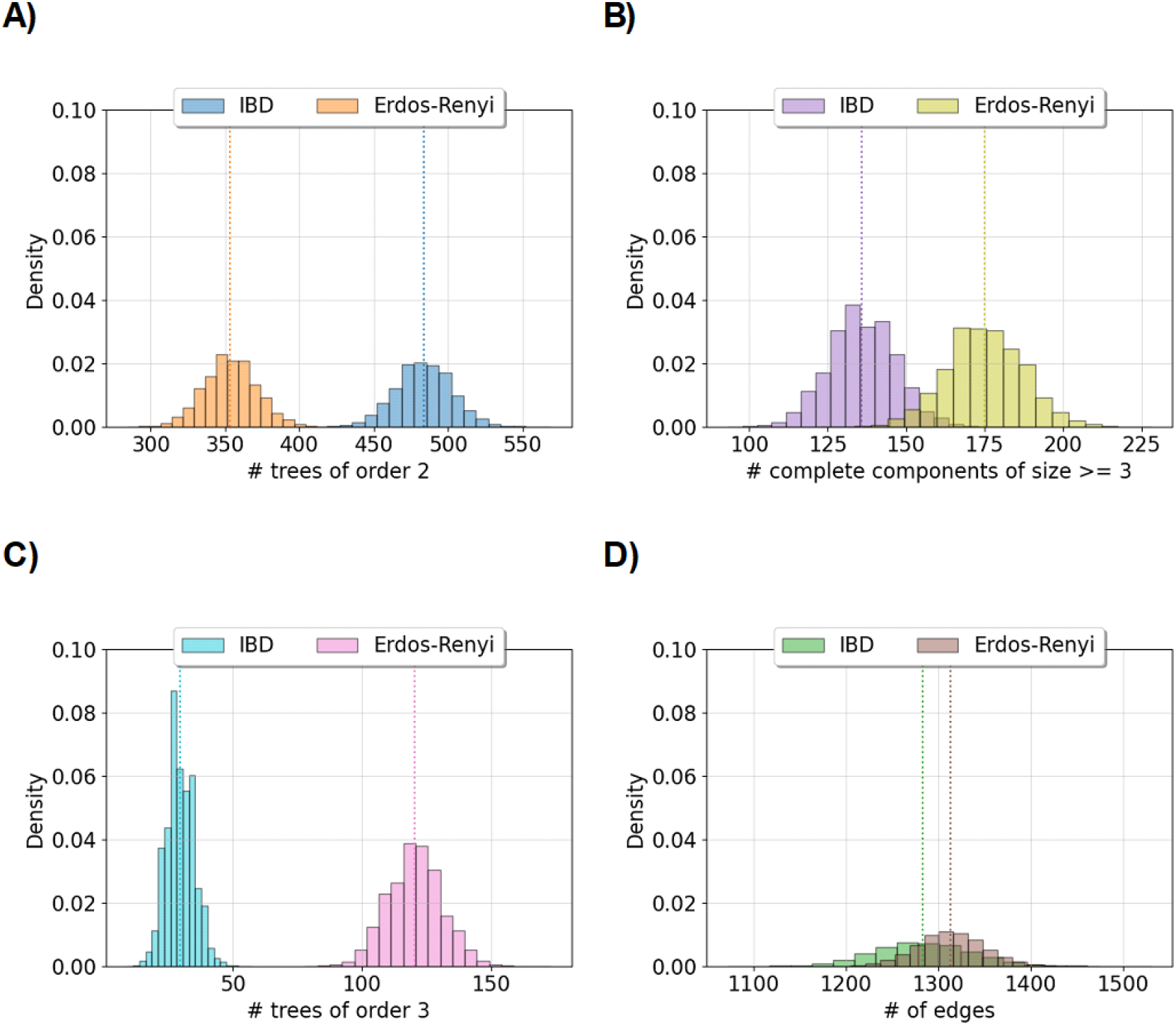
Comparing features between IBD and Erdős-Rényi graphs. Histograms compare the density of graph features between IBD and Erdős-Rényi graphs. Each histogram summarizes the results of one hundred and twenty-five thousand simulations. A) and C) show the number of trees of order 2 and 3, respectively. B) shows the number of complete components with more than three nodes. D) shows the total number of edges. The legends give color assigned to the IBD and Erdős-Rényi graphs. IBD graphs are simulated using the constant one hundred thousand diploid individuals’ demography and the 0.03 Morgans length threshold. Erdős-Rényi graphs are simulated using the same success probability as in the IBD graph. The sample size is two thousand diploid individuals. Vertical lines show the means.

#### 5.3.2. Flexible demographic scenarios

Figure S8 shows that the apparent normality of some graph features extends to the three phases of exponential growth and population bottleneck demographic scenarios. Table S1 reports that the proportions of rejected hypothesis tests for numbers of trees of order 2 are close to 0.05 for both demographic scenarios. We also cannot reject normality for the number of trees of order 3 and the number of connected components of degree 3 or more in some simulations of the three phases of exponential growth scenario. These results indicate that the limiting distributional behaviors of graph features in detectable IBD graphs around a locus can be similar for large constant populations and demographic scenarios with large recent population sizes.

#### 5.3.3. The impact of strong positive selection

Strong directional selection increases the detectable IBD rate around a locus [50] and the probability of IBD alleles [1], but less is known about how this phenomenon alters the feature distributions of detectable IBD graphs. In a hard selective sweep, a single allele increases in frequency at a rate of change that depends on a selection coefficient [14, 20, 25, 54]. The selection coefficient parameterizes the advantage that the sweeping allele has relative to alternative alleles, in so much as the gradient of the allele frequency trajectory is larger when the selection coefficient is larger.

We conduct more simulations of detectable IBD graphs for selection coefficients between 0.01 and 0.04 and the three phases of exponential growth and population bottleneck scenarios. Tables S2 and S3 demonstrate multiple trends as the selection coefficient increases. The apparent normality of the number of trees of order 2 does not noticeably change as we change the selection coefficient. Compared to our simulations with no selection, we reject normality less often for the number of trees of order 3 and the number of complete components of order 3 or more. It may be that the distributional behaviors of these small degree connected components become more apparent under the selection models with more detectable IBD segments. The main effect of strong positive selection appears to be the growth of the largest detectable IBD cluster that includes haplotypes with a beneficial allele. This idea is a major motivation for the suite of methods developed in Temple et al. [50].

## 6. Discussion

In this article, we leverage ideas from coalescent theory and haplotype sharing to develop statistical theory and motivate methodology in IBD-based inference. Most notably, we prove a central limit theorem for the detectable IBD rate around a locus whose regularity conditions have intuitive interpretations in population genetics. The sample size squared must be large enough such that there are many IBD segments long enough to be accurately detected by existing methods [5, 21, 44, 56]. The population size must be large enough that there are few to no large IBD clusters about a locus.

The conceptual framework for these conditions involves envisioning a coalescent tree with long internal branches, but numerous coalescent events occur near the leaves. The internal branches are long because of the large population size, and there are numerous coalescent events near the leaves because of the large sample size. The large Morgans threshold further decreases the probability of a detectable IBD segment and the correlations between IBD segment indicators.

The techniques we use might be useful to other studies involving coalescent *and* recombination processes. For instance, to generalize our main central limit theorem, we take a formulaic approach. First, we derive covariances for a finite set of classes. Second, we count the number of covariance terms of each class that occur in the total covariance of the sample mean. Third, we determine a “little-o” condition such that the sum of covariances of one specific class is asymptotically equivalent to the sum of covariances of all the other classes. We use a particular central limit theorem for dependent data [11, 12], which is derived using Stein’s moment-based method—a more general technique to demonstrate weak convergences to Gaussian or non-Gaussian random variables [29, 40, 46, 47].

One counterexample in which our particular proof strategy does not work concerns the density of recent coalescent events (DRC) [38]. This test statistic is the sample mean of indicators if a haplotype pair has a common ancestor within a given time threshold ^3^. For constant population size *N*, each covariance type is the survival function of a hypoexponential random variable. The main terms in all these survival functions take the form exp (−*C* · *N*), where *C* is a constant. However, the combinatorics are the same as those in our IBD-based statistics. As a result, the sum of covariances of one specific class is *not* asymptotically equivalent to the sum of covariances of all the other classes. This observation points out the role that integrating over *shared recombinations* plays in reducing the covariance. We employ simulation to evaluate the assumptions and validity of our central limit theorem. Consistent with our conditions, we reject a null hypothesis of normality less often as sample size and scaled population size increase. In practice, we find that non-normality is typical in finite samples. We indicate that nonnegligible covariance may come from the accumulation of IBD clusters. Based on the tail behavior of simulated distributions, we expect that a one-sample *z* test for excess IBD rates may inflate the number of false positives.

Our regularity conditions concern a balance between sample size and scaled population size that is unlikely to hold in practical settings. In our experiments, we observe neither a trend between sample size and the proportion of rejected tests nor between sample size and the relative upper tail probability. We advocate that the collected sample size should always be as large as is feasible and that the smallest Morgans length threshold for which IBD segment detection is accurate should be chosen.

Our theoretical results and simulation studies support ongoing methodological developments based on IBD segments. Existing genome-wide scans for excess IBD rates [5, 50] or differences in IBD rates between groups [7] lack formal or exact hypothesis testing frameworks. Motivated in part by this work, Temple [48] controls the family-wise error rate (FWER) in their selection scan by modelling the IBD rate process as an Ornstein-Uhlenbeck process, thereby assuming that the IBD rate is normally distributed at any given spatial position. Consistent with this work, they show anti-conservative control of FWER. Combining an FWER control technique [19, 45] with our multivariate central limit theorem, we indicate that a modification of the Temple [48] method may apply to a test for equality of detectable IBD rates in case-control studies. In these examples and others [13, 22, 32] from statistical and population genetics, assuming reasonable asymptotic models is often vital when adjusting for many correlated tests.

## Data and code availability

We use the Python package https://github.com/sdtemple/isweep for all simulation studies. This software is freely available under the open-source CC0 1.0 Universal License.

## Acknowledgements

S.D.T. acknowledges funding support from the US National Defense Science and Engineering Graduate Fellowship, the US National Institute of Health T32 GM081062 Pre-doctoral Training Grant in Statistical Genetics, and Schmidt Sciences LLC. Computational resources were supported in part by the National Human Genome Research Institute of the National Institutes of Health under award number HG005701 and the Department of Statistics at the University of Washington. We thank Sharon R. Browning for helpful discussions.

## Author contributions

S.D.T. proposed the study, planned the study, wrote the software, conducted the simulation studies, and wrote the manuscript. Both authors worked on the theoretical results and contributed to editing the manuscript.

## A.1. Derivations of theoretical results

### A.1.1. Theorem 3.1 and its extensions

#### Lemma A.1.

𝔼_2_[*X*_*a,b*_] → 0 *uniformly as Nw* → ∞.

*Proof*. Let *f* (*N*) = (*Nw*)^−1^, and recall that *E*_2_[*X*_*a,b*_] = (2*Nw* + 1)^−1^. If *Nw >* (1*/ε* − 1)*/*2, then |*f* (*N*) − 0| < *ε*. Choose integer *M* such that *Mw* ≥ (1*/ε* − 1)*/*2. Thus, for *ε* > 0, there exists *M* such that |*f* (*N*)| = (2*Nw* + 1)^−1^ < *ε* for all *N* ≥ *M*.

#### Lemma A.2.

*Let X* ∼ *Bernoulli*(*q*) *and q* ∈ (0, 1). 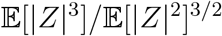 *is bounded above where Z* = *X* − 𝔼 [*X*].

*Proof*.

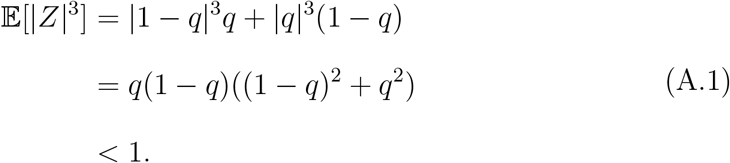

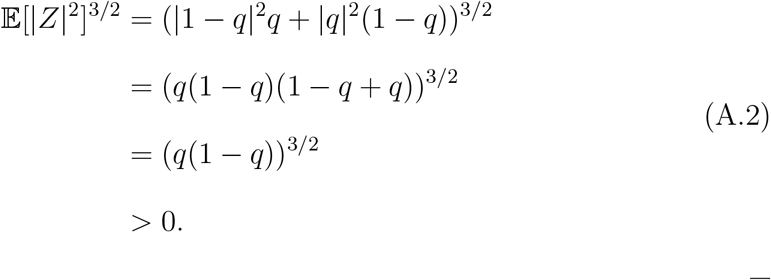

#### Lemma A.3.

*Cov*_3_(*Z*_*a,b*_, *Z*_*a,c*_) ≡ *Cov*_3_(*X*_*a,b*_, *X*_*a,c*_) = *O*((*Nw*)^−2^).

*Proof*. Up to reordering three sample haplotypes, there is one possible bifurcating tree (Figure S9). Sample haplotypes *a* and *b* coalesce to a common ancestor, and their common ancestor coalesces to a common ancestor with sample haplotype *c*. We integrate over coalescent time and haplotype segment lengths to bound the covariance.

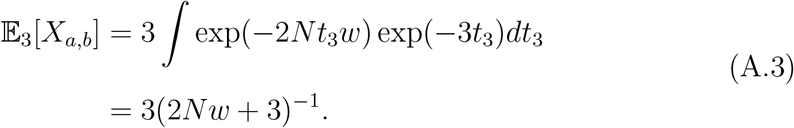

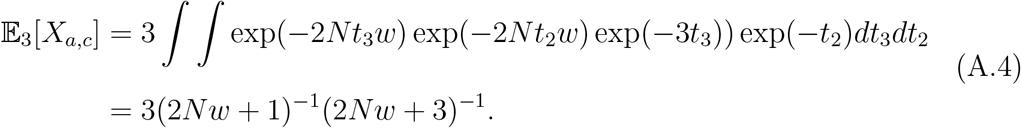

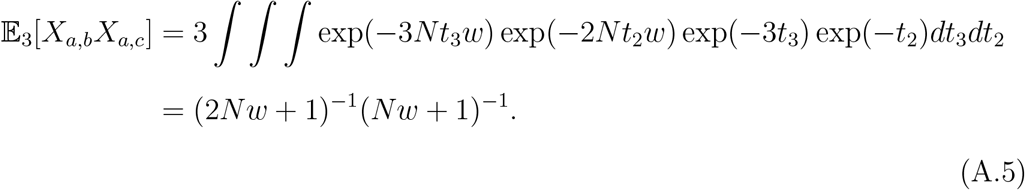

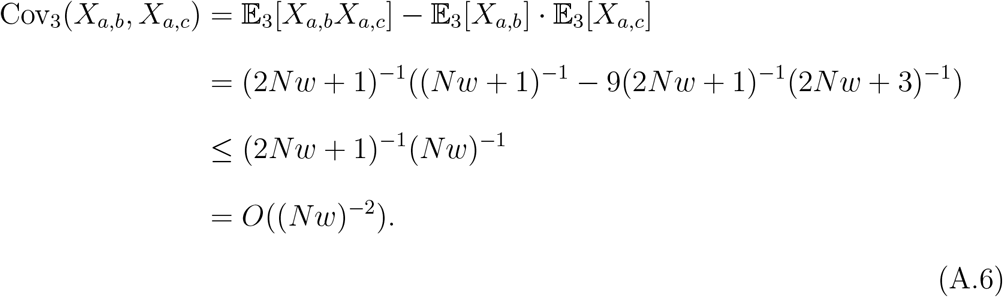

#### Lemma A.4.

*Cov*_4_(*Z*_*a,b*_, *Z*_*c,d*_) ≡ *Cov*_4_(*X*_*a,b*_, *X*_*c,d*_) = *O* ((*Nw*)^−3^).

*Proof*. Up to reordering four sample haplotypes, there are two possible bifurcating trees (Figure S10). The first tree is as follows: sample haplotypes *a* and *b* coalesce to a common ancestor, then sample haplotypes *c* and *d* coalesce to a common ancestor, and finally those common ancestors coalesce. The covariance of *X*_*a,b*_ and *X*_*c,d*_ is zero because of independent meioses. We focus instead on the covariance of *X*_*a,c*_ and *X*_*b,d*_. We integrate over coalescent time and haplotype segment lengths to bound the covariance.

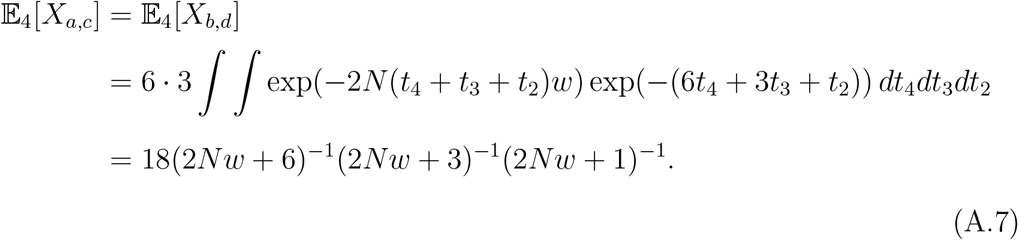

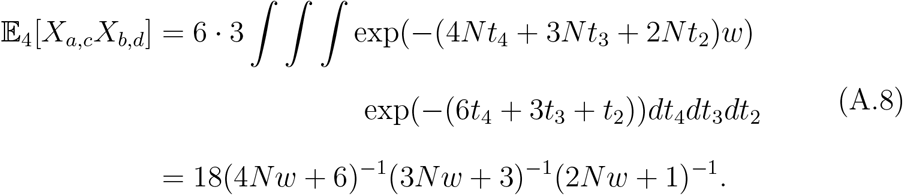

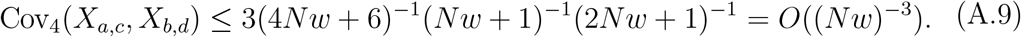

The second tree is as follows: *a* and *b* coalesce to a common ancestor, then their common ancestor coalesces with *c*, and finally, the common ancestor of *a, b*, and *c* coalesces with *d*. It is easy to verify that 𝔼_4_[*X*_*a,c*_*X*_*b,d*_] is the exact same as in Equation A.8. Next,

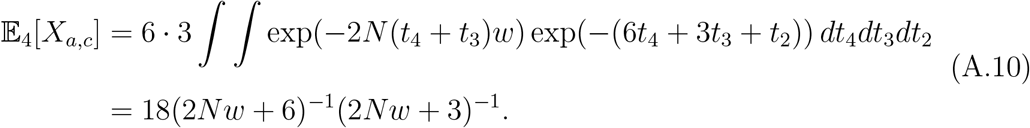

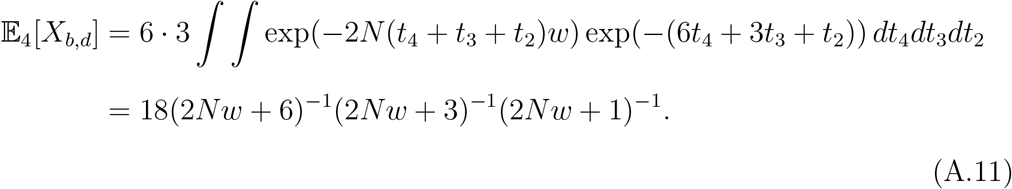

Because Equations A.10 and A.11 are nonnegative, the marginal covariance upper bound is the same as in Equation A.9.

#### Lemma A.5.

*The following are true*

- 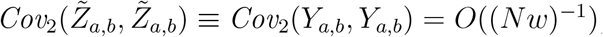
- 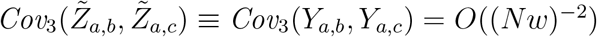
- 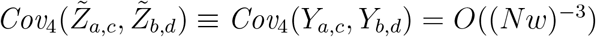

*Proof*. We take the same approach as in Lemmas A.3 and A.4, except the survival function is that of an Erlang random variable with shape parameter 2.

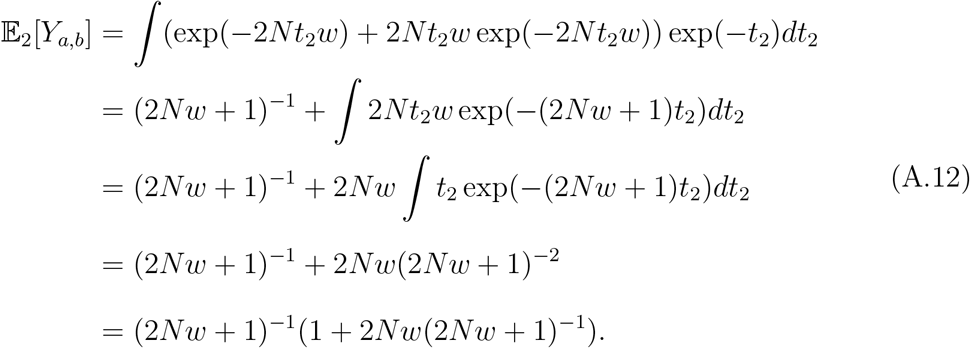

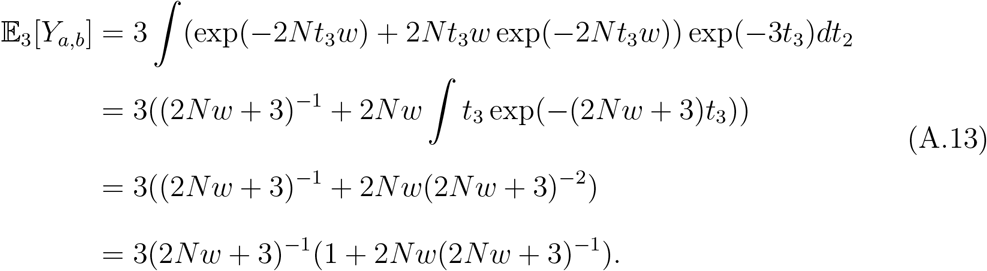

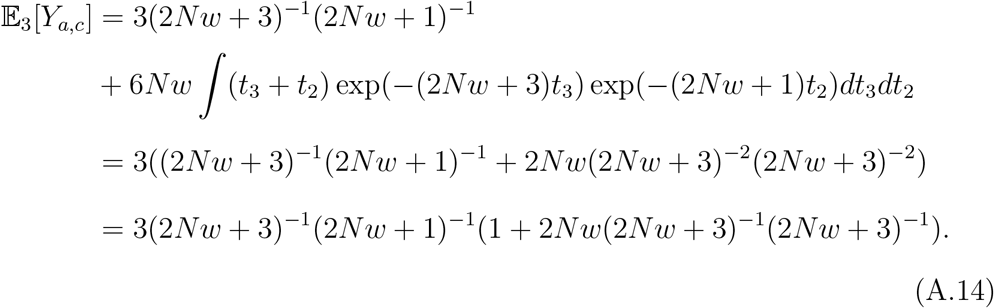

From Equations A.12, A.13, and A.14, the pattern emerges that the effect of the convolution of crossover points is to multiply *O*(1) terms to the marginal expected values in Equation 4 and Lemmas A.3 and A.4.

Calculating 𝔼_3_[*Y*_*a,b*_*Y*_*a,c*_] is more involved. Up to reordering three sample hap-lotypes, we consider sample haplotypes *a* and *c* that coalesce at the most recent common ancestor of *a, b*, and *c*. Then, 𝔼_3_[*Y*_*a,c*_] ≥ 𝔼_3_[*Y*_*a,b*_*Y*_*a,c*_], and

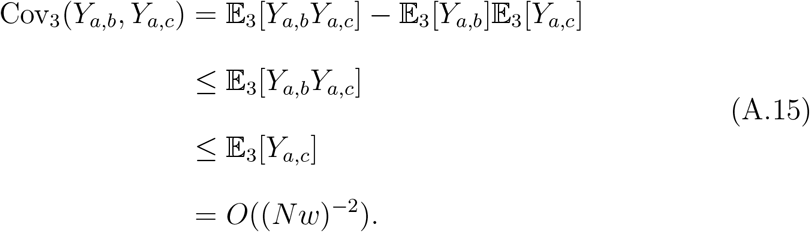

Using the same techniques, it is easy to calculate 𝔼_4_[*Y*_*a,c*_] and 𝔼_4_[*Y*_*b,d*_] for the two different tree shapes and derive the *O* ((*Nw*)^−3^) bound for Cov_4_(*Y*_*a,c*_, *Y*_*b,d*_).

#### Lemma A.6.

*For a sample of three haplotypes a, b, and c, when* 𝔼_2_[*X*_*a,c*_] *<* 1*/*2, *the conditional expectation* 𝔼 [*Z*_*a,c*_ *×* **Z**_−*a,c*_|**Z**_−*a,c*_] ≱0 *for all* **Z**_−*a,c*_.

*Proof*. Define *q* =: 𝔼_2_[*X*_*a,c*_], and fix **X**_−*a,c*_ = 1.

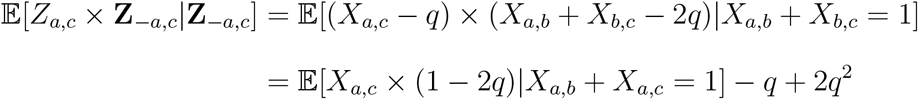

Because of IBD transitivity, *X*_*a,c*_ = 0 with probability 1. Then, the equation simplifies to −*q*(1 − 2*q*) < 0.

### A.1.2. Multi-way IBD segments

*Proof of Theorem 4.2*. We give the general argument for 3-way IBD segment indicators. To begin, we calculate bounds on the relevant integrals 𝔼_*κ*_[·],…, 𝔼_2*κ*_[·]. Recall that E_*κ*_ is the expected value with respect to a coalescent tree of *κ* haplo-types.

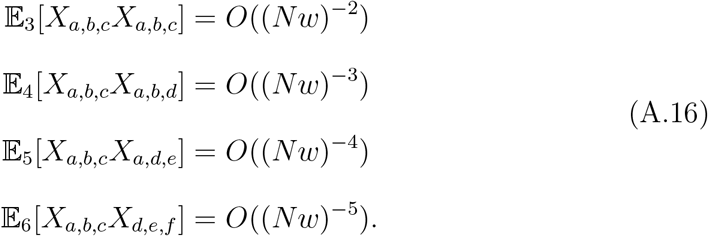

These are also the covariance bounds because 𝔼_3_[*X*_*a,b,c*_*X*_*a,b,c*_] ≥ 0 and 𝔼_3_[*X*_*a,b,c*_*X*_*a,b,c*_] ≥ 𝔼_3_[*X*_*a,b,c*_]^2^ and so on for the other 𝔼_*κ*_ relations.

Next, we take sums over these covariance bounds and substitute in the *n* =*o*(*Nw*) condition.

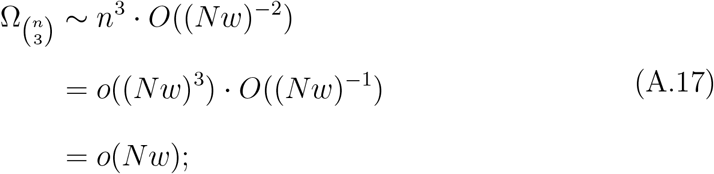

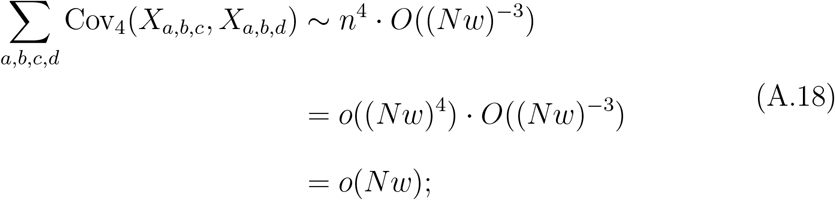

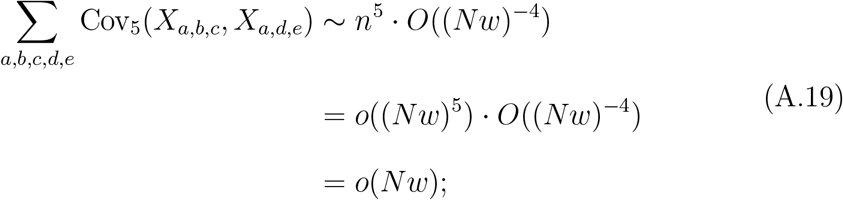

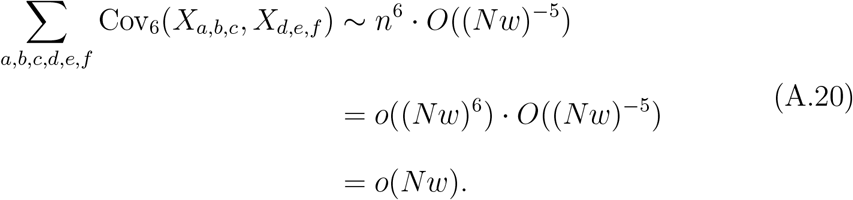

The covariance within IBD segment indicators 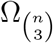 controls the sum of covariances Σ _(*a,b,c)*≠ (*d,e,f*)_ Cov(*X*_*a,b,c*_, *X*_*d,e,f*_). Using the bounding argument in Equation A.15, the result extends to IBD segment indicators around a focal location.

### A.1.3. Multivariate IBD rates

*Proof of Theorem 4.3*. To use Corollary 1 from Chandrasekhar et al. [12] in multiple dimensions, we now require defining their notion of an affinity set. These are subsets 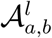 containing the haplotype pair *a* and *b* from sample *l* such that 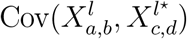 is high if the haplotype pair *c* and *d* from sample *l*^⋆^ are in the affinity set and low if they are not. We consider the singleton affinity sets 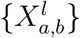. (We remark that singletons are the affinity sets we use in all of our proofs for the one-dimensional results.)

We use the example of two sample means to concretely calculate covariances. Let Ω_2*×*2_ be the covariance matrix such that

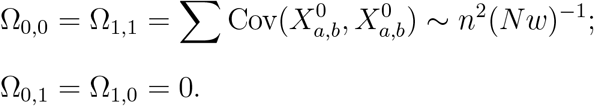

Ω_0,1_ and Ω_1,0_ concern the sum of covariances of IBD segment indicators within affinity sets but in different samples, which is zero because the affinity set of a haplotype pair in one sample includes no haplotype pairs in a different sample.

The term that controls the sum of covariances across affinity sets is the Frobenius norm ||Ω_2*×*2_||_*F*_. We calculate this norm as

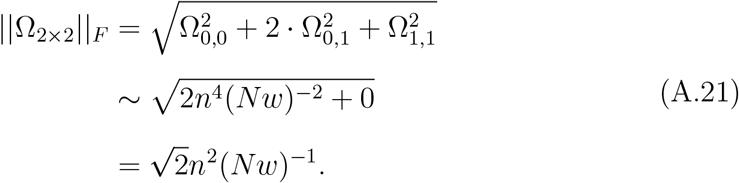

Under the condition *n* = *o*(*Nw*), Equation A.21 is *o*(*Nw*), and under the condition *Nw* = *o*(*n*^2^), the variance term ||Ω_2*×*2_||_*F*_ tends to infinity.

The first condition from Corollary 1 in Chandrasekhar et al. [12] is

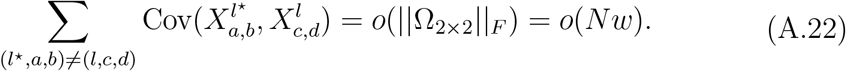

We compute the sums of covariances of IBD segment indicator types {(*a, b*), (*a, e*)} and {(*a, b*), (*c, d*)}, where *a, b*, and *e* are haplotypes in one sample and *c* and *d* are haplotypes in the other sample. By the same calculations as in the previous proofs, these sums are asymptotically equivalent to *n*^3^(*Nw*)^−2^ = *o*(*Nw*) and *n*^4^(*Nw*)^−3^ = *o*(*Nw*).

Since the column vector is now multi-dimensional, we must also show that

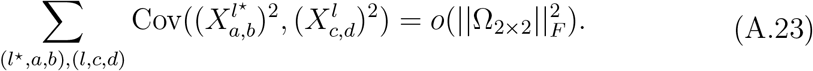

This calculation is simplified as

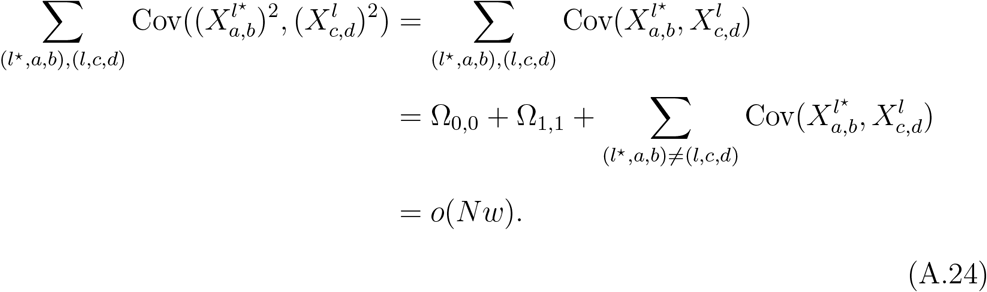

We get the general result by extending these calculations for sums and norms over covariances of two samples to those of 𝓁samples. The term in Equation A.22 involves sums of covariances of 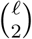 pairs of samples. This term is why we require the bound on 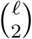, because in Equation A.21 we have the multiplicative factor 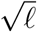.

Using the bounding argument in Equation A.15, the result extends to IBD segment indicators around a focal location.

## A.2. Verifying an assumption of the central limit theorem

We take a Monte Carlo approach to examine the conditional expectation assumption 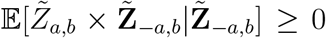 for all 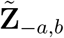 because 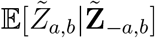 is analytically intractable. Namely, by replacing the expected value 𝔼[*Y*_*a,b*_|**Y**_−*a,b*_] with an average over a large number of simulations, we assess if 𝔼 [*Y*_*a,b*_|**Y**_−*a,b*_] ≥ 𝔼[*Y*_*a,b*_] when 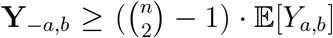 and vice versa that 𝔼 [*Y*_*a,b*_|**Y**_−*a,b*_] ≤ 𝔼 [*Y*_*a,b*_] when 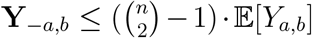. (Recall that *Z*_*a,b*_ is the binary random variable *Y*_*a,b*_ after mean-centering.) The intuition is that if the observed sum **Y**_−*a,b*_ is larger than the expected sum 𝔼 [**Y**_−*a,b*_] then the held out *Y*_*a,b*_ is more likely to be 1 than it would be if the observed sum equaled the expected sum.

We run the Temple et al. [49] algorithm one hundred and twenty million times, recording the value of *Y*_*a,b*_ and the sum **Y**_−*a,b*_ for some fixed haplotype pair *a* and *b*. Then, we calculate the difference between the empirical average 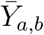 and 𝔼[*Y*_*a,b*_], stratified into eight quantile bins depending on the sum **Y**_−*a,b*_. The sample sizes are limited to two to four hundred diploid individuals to keep runtime modest.

Figure S11 shows the results of this simulation study. For each bin, the average count is less than and greater than 𝔼 [*Y*_*a,b*_] when the sum **Y**_−*a,b*_ is less than and greater than 𝔼 [**Y**_−*a,b*_], respectively. This trend is especially apparent for **Y**_−*a,b*_ far from the mean IBD count 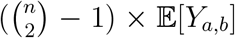. These findings provide empirical evidence that the theorem assumption may be true for moderate to large sample sizes.

**Figure S1:**
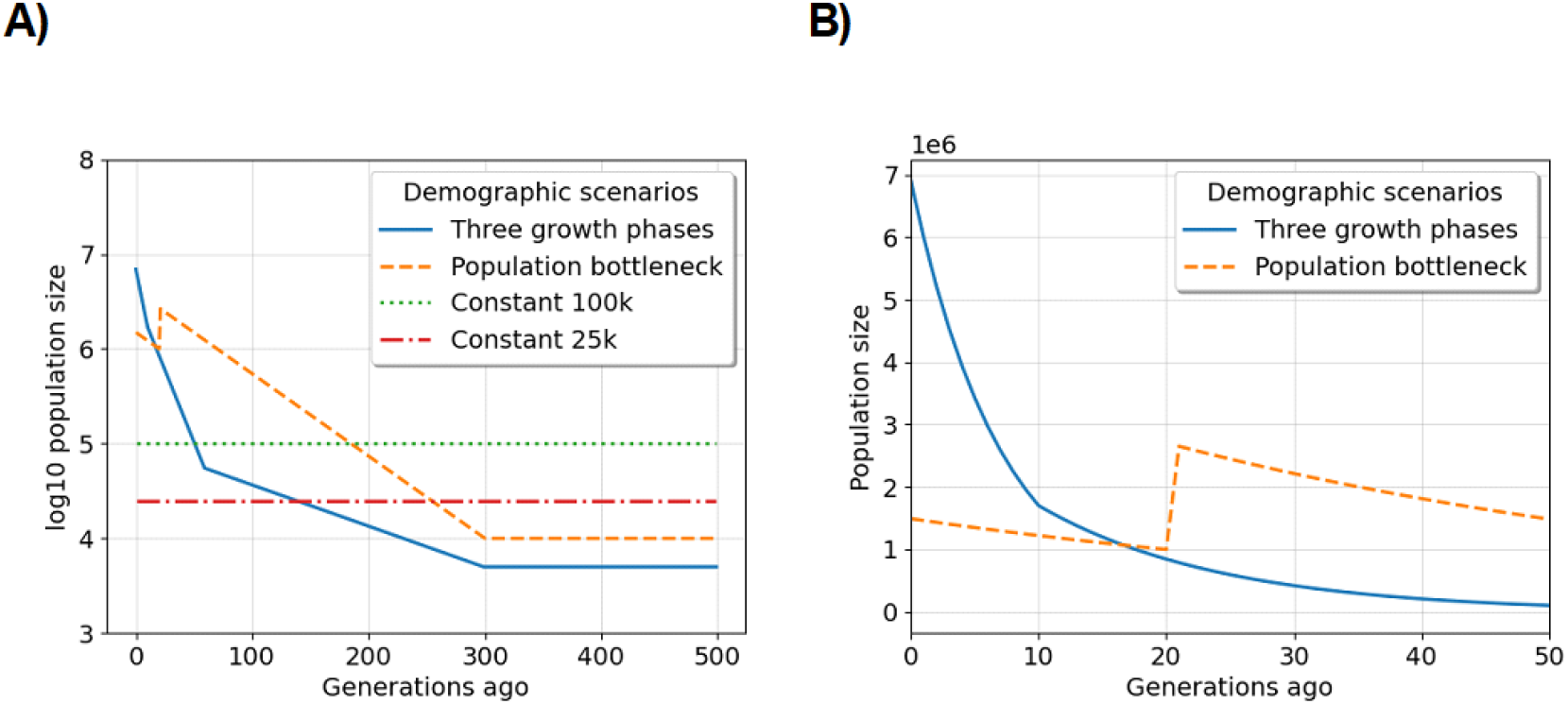
Demographic scenarios we consider in simulation studies: A) coalescent time in generations ago by the log 10 population size, and B) the most recent fifty generations by population size for examples of exponential growth. The legends specify the color and line style for each scenario. As opposed to coalescent time used in the main text, we describe the scenarios forward in time here. Three phases of exponential growth: a population of ancestral size five thousand diploids increases exponentially each generation at rates one, seven, and fifteen percent starting three hundred, sixty, and ten generations ago. Population bottleneck: a population of ancestral size ten thousand diploids increases exponentially each generation at a rate of two percent starting three hundred generations ago. Otherwise, the demographic scenarios we explore here are populations of constant size twenty-five and one hundred diploids.

**Figure S2:**
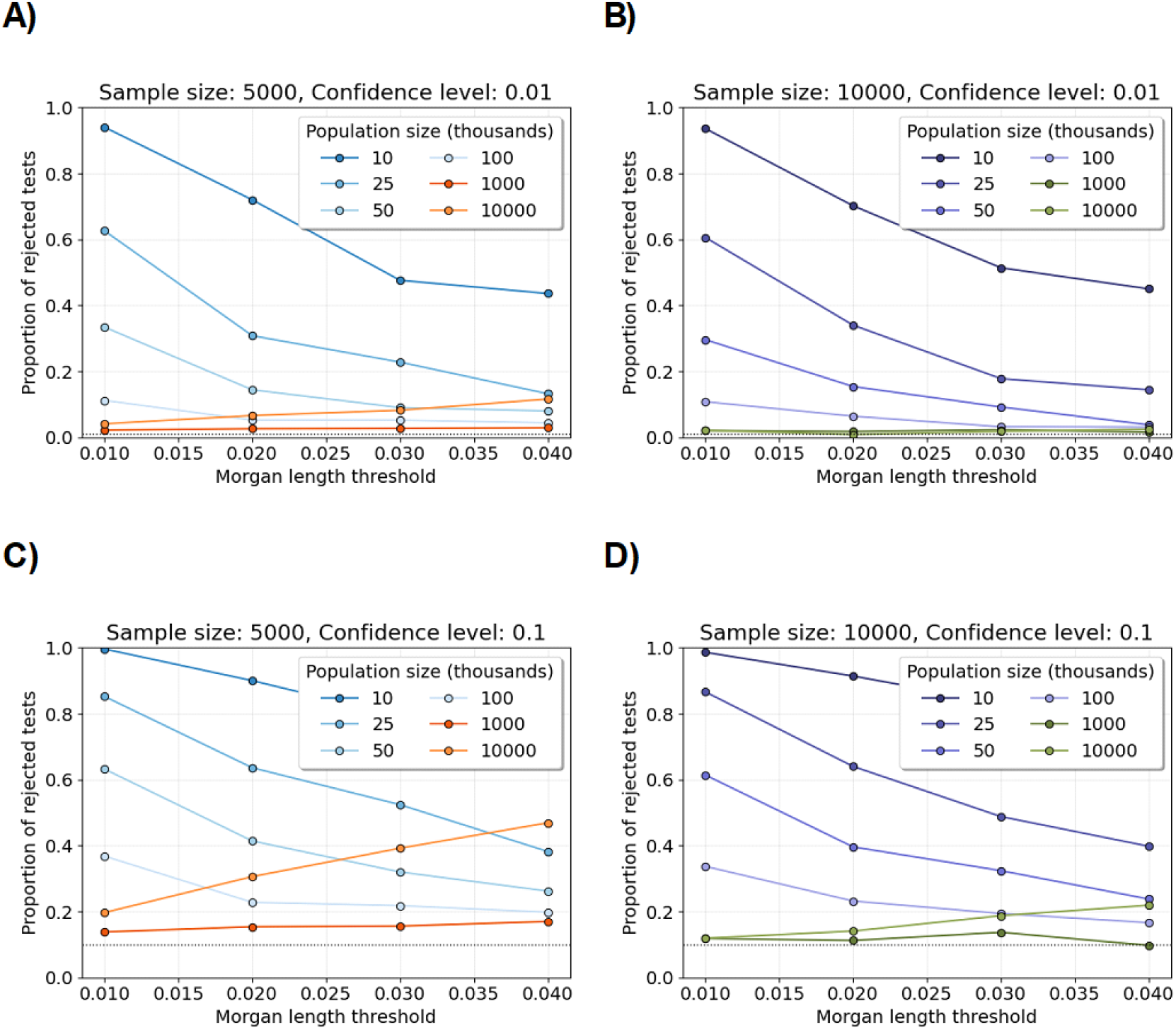
Shapiro-Wilk tests for varying population sizes and significance levels. Line plots show the proportions of Shapiro-Wilk tests rejected at significance levels A,B) 0.01 and C,D) 0.1 (y-axis) for varying population size and fixed sample size. Each proportion is computed over five hundred tests. Each test is based on one thousand simulations of the number of identity-by-descent lengths longer than a specified Morgans length threshold (x-axis). A,C) The sample size is five thousand diploid individuals. B,D) The sample size is ten thousand diploid individuals. The legends assign colors to different population sizes. The horizontal dotted lines are significance levels.

**Figure S3:**
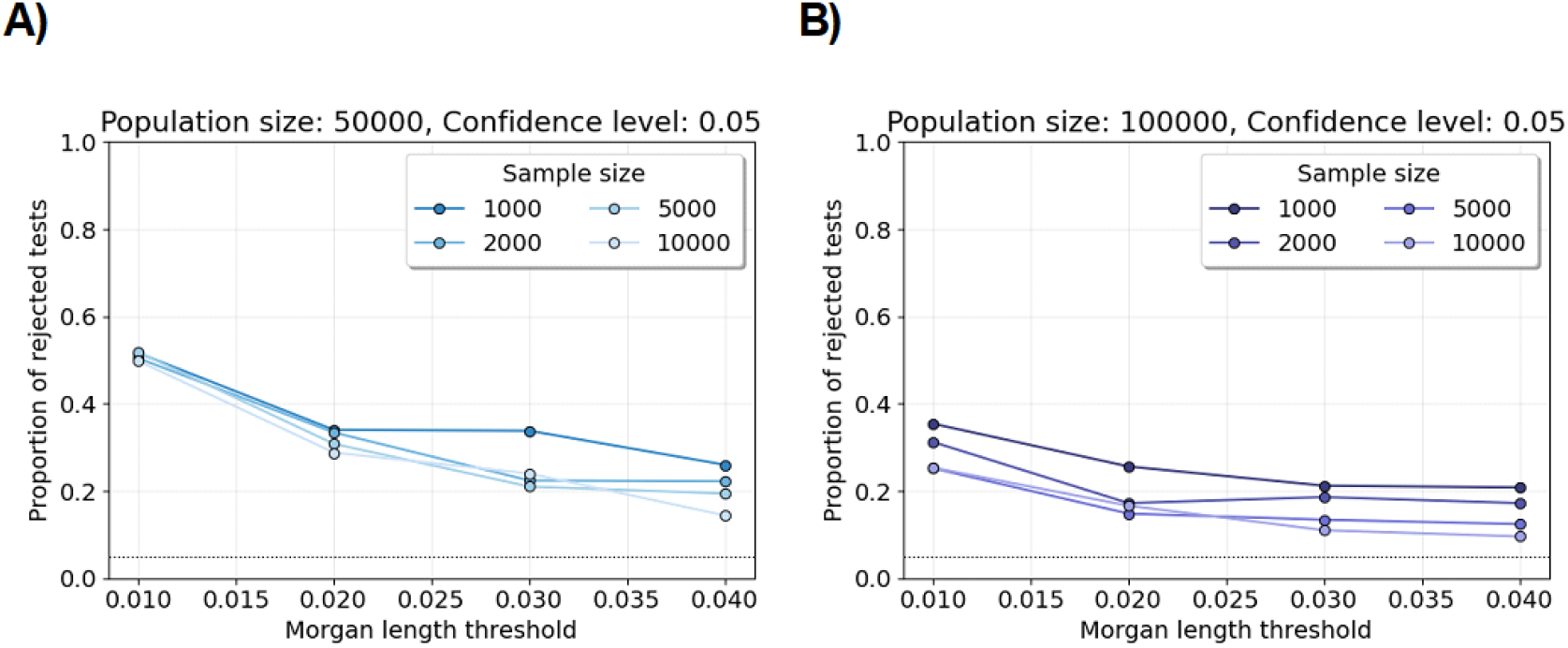
Shapiro-Wilk tests for varying sample sizes. Line plots show the proportions of Shapiro-Wilk tests rejected at the significance level 0.05 (y-axis) for varying sample size and fixed population size. Each proportion is computed over five hundred tests. Each test is based on one thousand simulations of the number of identity-by-descent lengths longer than a specified Morgans length threshold (x-axis). A) The population size is fifty thousand diploid individuals. B) The population size is one hundred thousand diploid individuals. The legends assign colors to different sample sizes. The horizontal dotted line is at 0.05.

**Figure S4:**
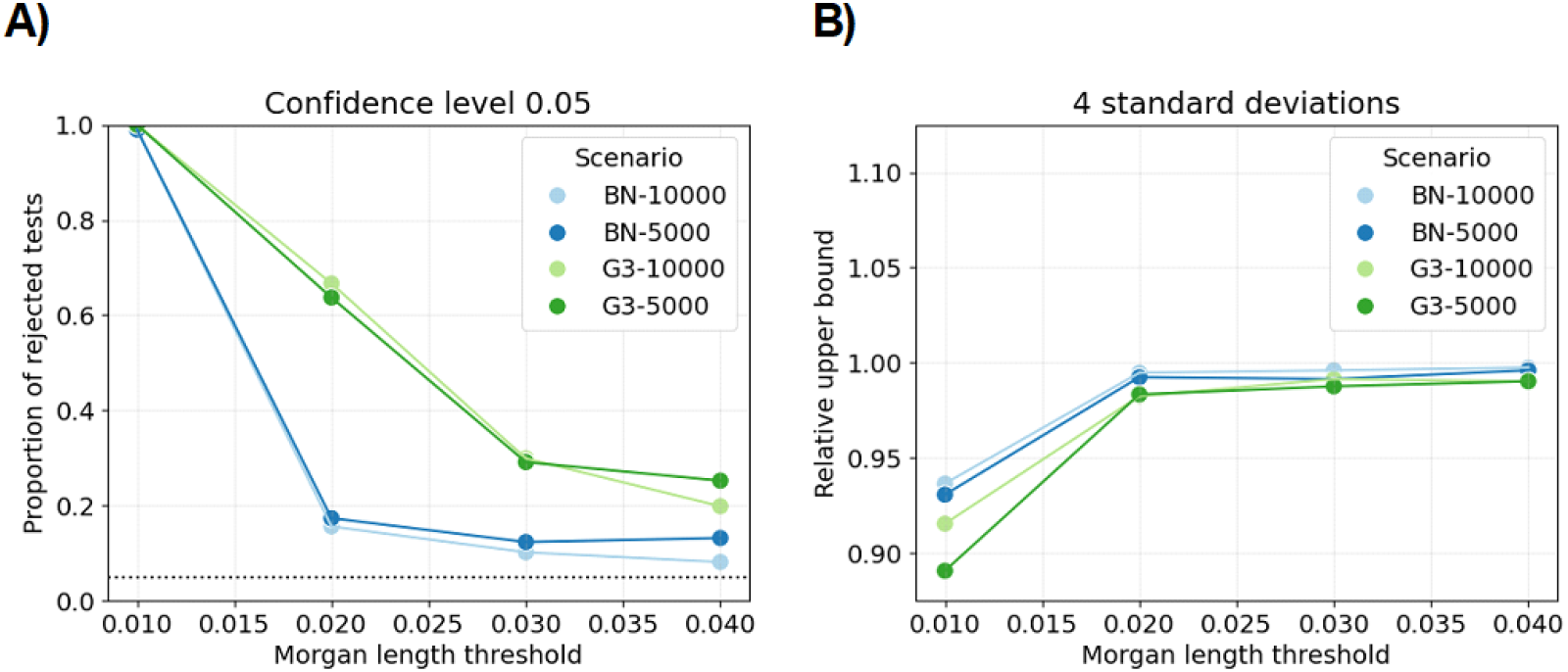
Shapiro-Wilk tests and relative upper tail bounds for complex demography scenarios. A) Line plots show the proportions of Shapiro-Wilk tests rejected at the sigificance level 0.05 (y-axis) for the population bottleneck (BN) or three phases of exponential growth (G3) demographic scenarios and sample sizes of five or ten thousand diploid individuals. Each proportion is computed over at least six hundred tests. Each test is based on one thousand simulations of the number of identity-by-descent lengths longer than a specified Morgans length threshold (x-axis). B) Line plots show the average mean plus four standard deviations divided by the 99.99683 percentile over two million simulations (y-axis). Plot designs are identical to Figures 2 and 3.

**Figure S5:**
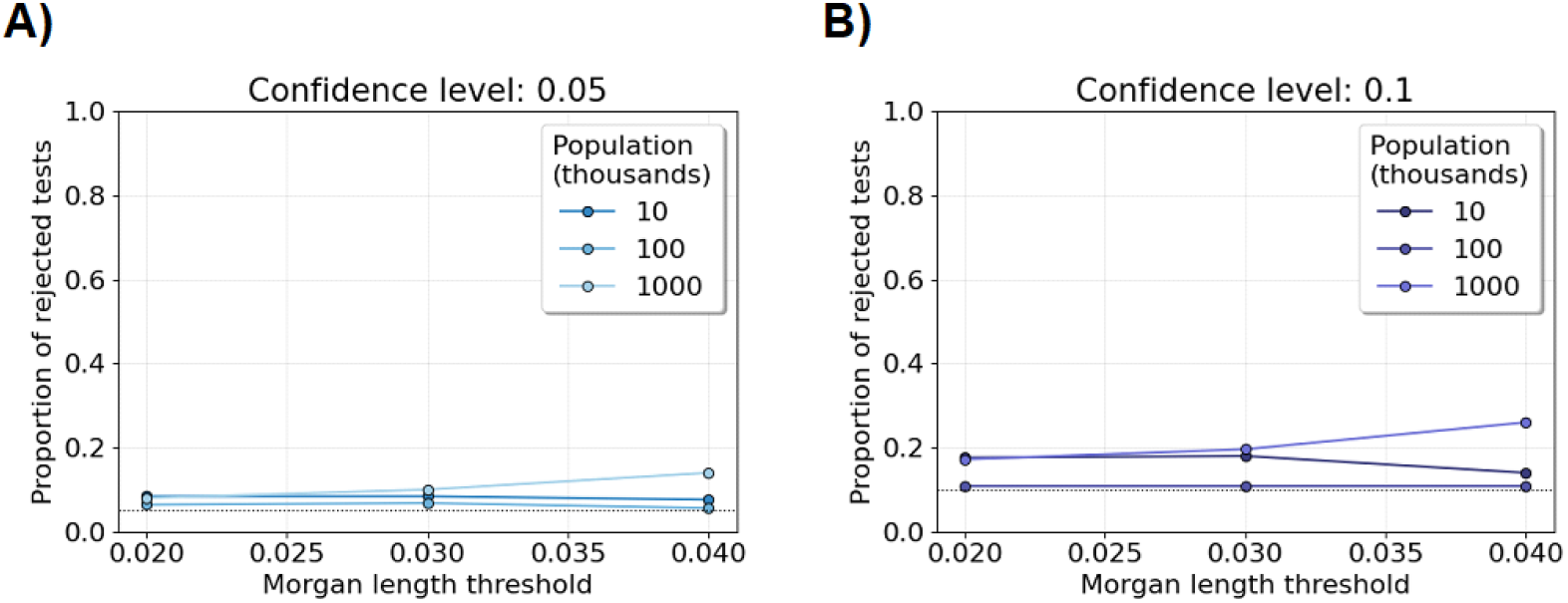
Shapiro-Wilk tests for difference in IBD rates between groups. Line plots show the proportions of Shapiro-Wilk tests rejected at the significance level 0.05 (y-axis) for increasing constant population sizes (in thousands). The sample size is five thousand diploid individuals. Each proportion is computed over two hundred and fifty tests. Each test is based on five hundred simulations of the difference between groups in IBD rates longer than a specified Morgans length threshold (x-axis). The significance threshold is either A) 0.05 or B) 0.10, shown as horizontal dotted black lines.

**Figure S6:**
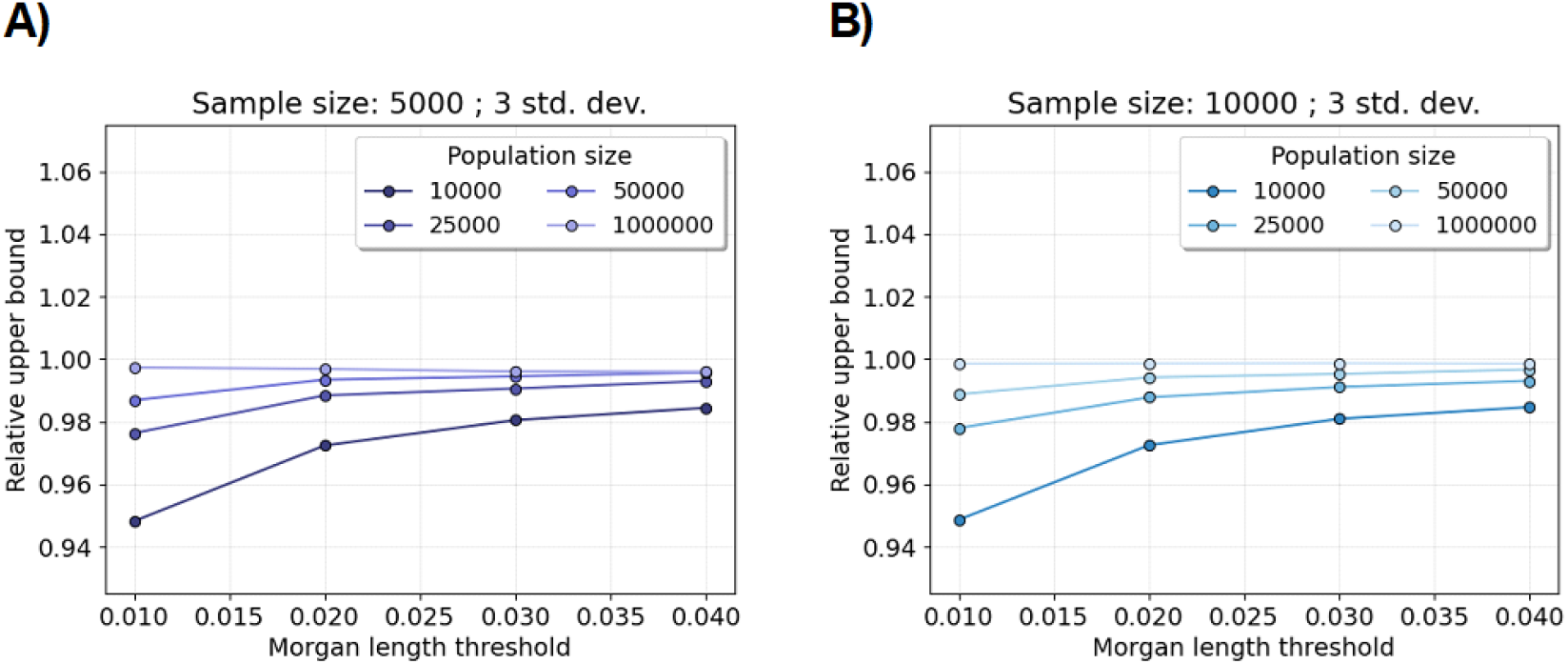
Relative upper bound for excess IBD scan. Line plots show the average mean plus three standard deviations divided by the 99.86501 percentile over two million simulations (y-axis). (The standard normal survival function of three is 0.9986501.) Each average relative upper bound is computed over one thousand tests. Each test is based on two thousand simulations of the number of identity-by-descent lengths longer than a specified Morgans length threshold (x-axis). A) The sample size is five thousand diploid individuals. B) The sample size is ten thousand diploid individuals. The legends assign colors to different constant population sizes.

**Figure S7:**
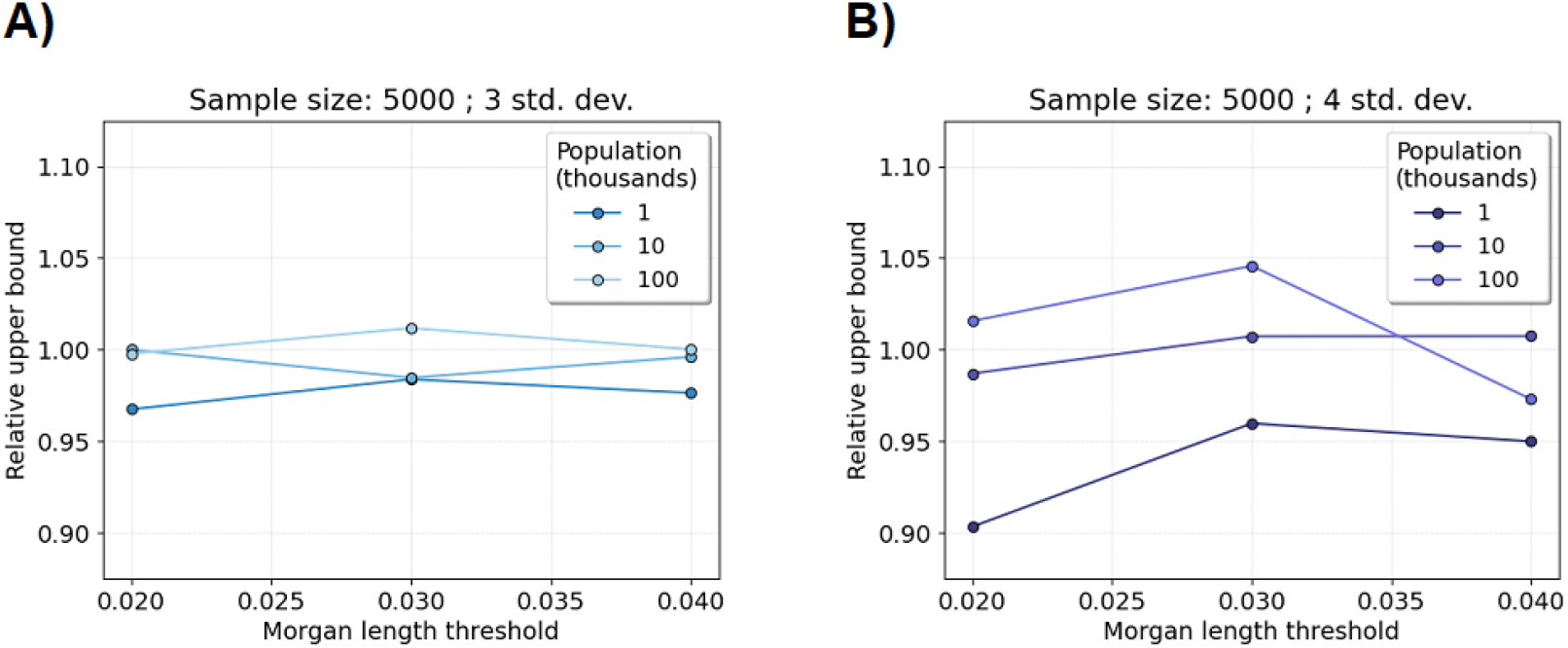
Relative upper bound for the difference in IBD rates test. Line plots show the average mean plus A) three or B) four standard deviations divided by the standard normal corresponding percentiles over one hundred and twenty-five thousand simulations (y-axis). Each average relative upper bound is computed over two hundred and fifty tests. Each test is based on five hundred simulations of the number of identity-by-descent lengths longer than a specified Morgans length threshold (x-axis). The sample size is five thousand diploid individuals. The legends assign colors to increasing constant population sizes (in thousands).

**Figure S8:**
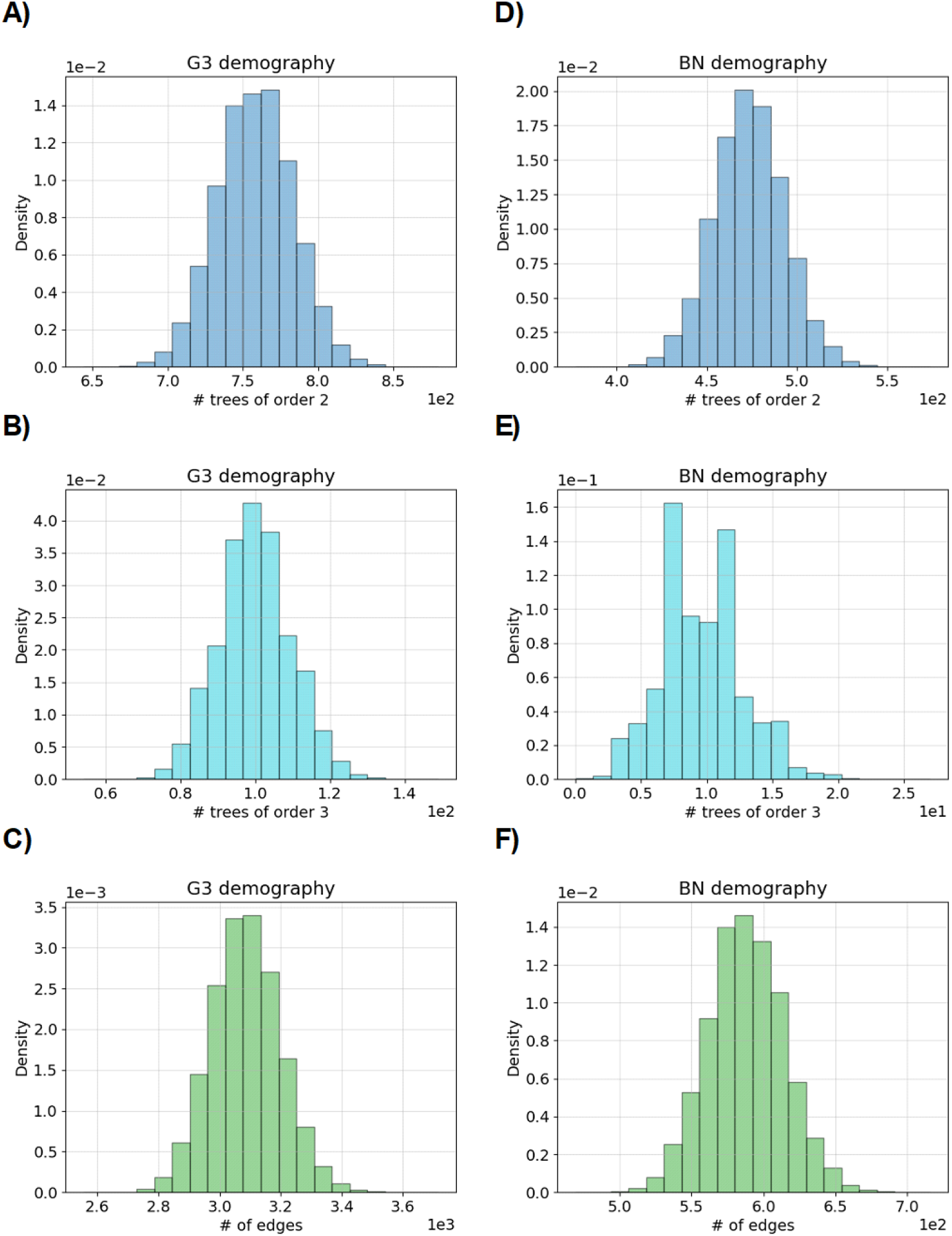
Comparing features between IBD graphs for complex demographic scenarios. Histograms show the density of IBD graph features between A-C) the three phases of exponential growth (G3) and D-F) the population bottleneck (BN) demographic scenarios. Each histogram is based on at least six hundred thousand simulations. A,D), B,D), and C,F) show the number of trees of order 2, the number of trees of order 3, and the total number of edges, respectively. The Morgans length threshold is 0.03. The sample size is five thousand diploid individuals.

**Figure S9:**
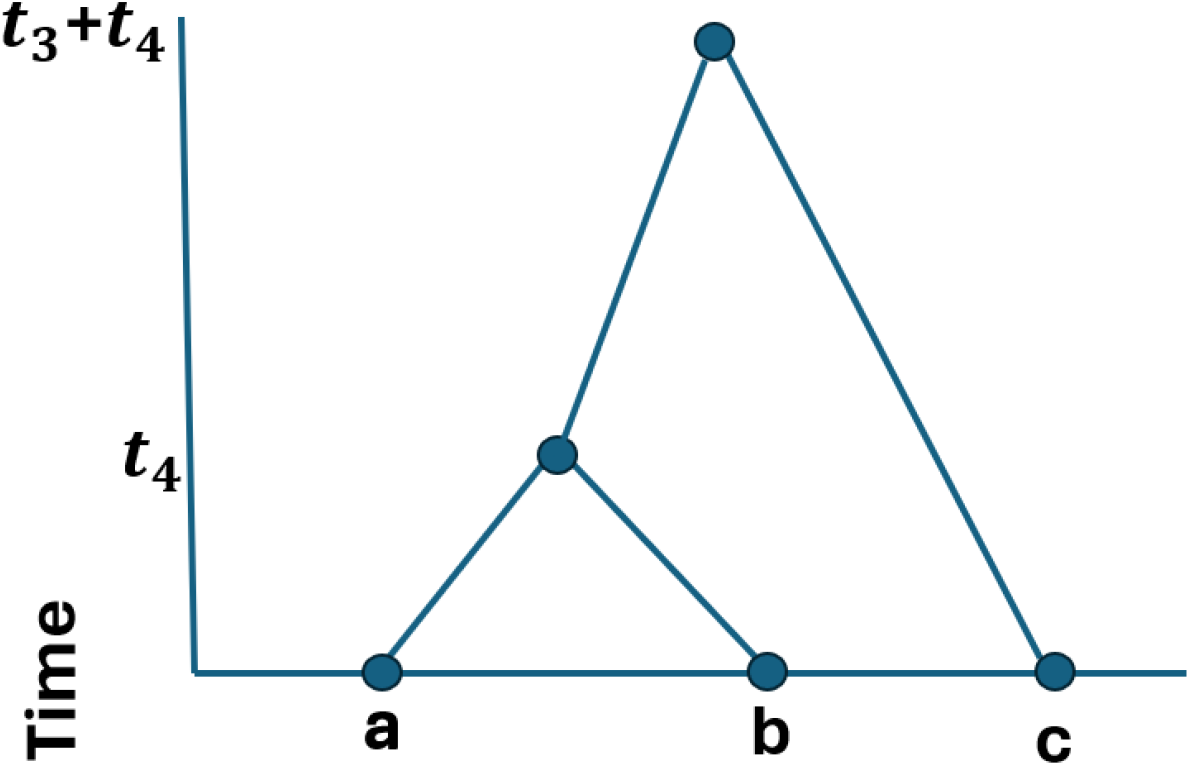
Illustration of the one possible coalescent tree used to calculate Cov_3_ terms in Appendix A.1.

**Figure S10:**
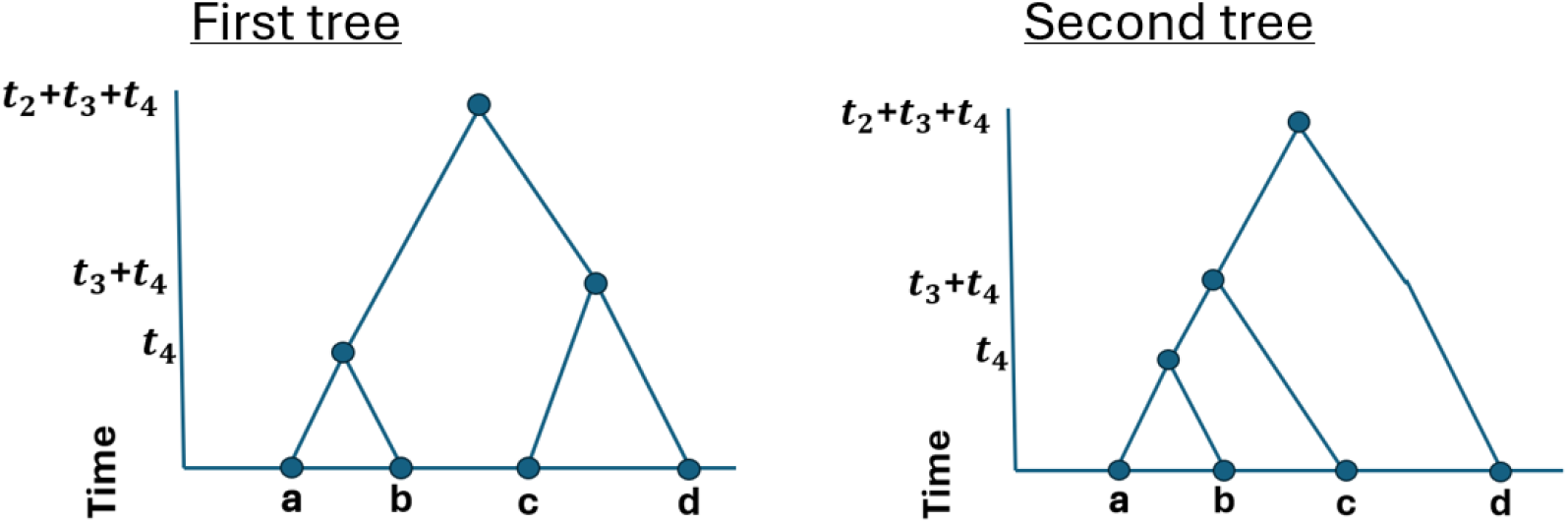
Illustration of the two possible coalescent trees used to calculate Cov_4_ terms in Appendix A.1.

**Figure S11:**
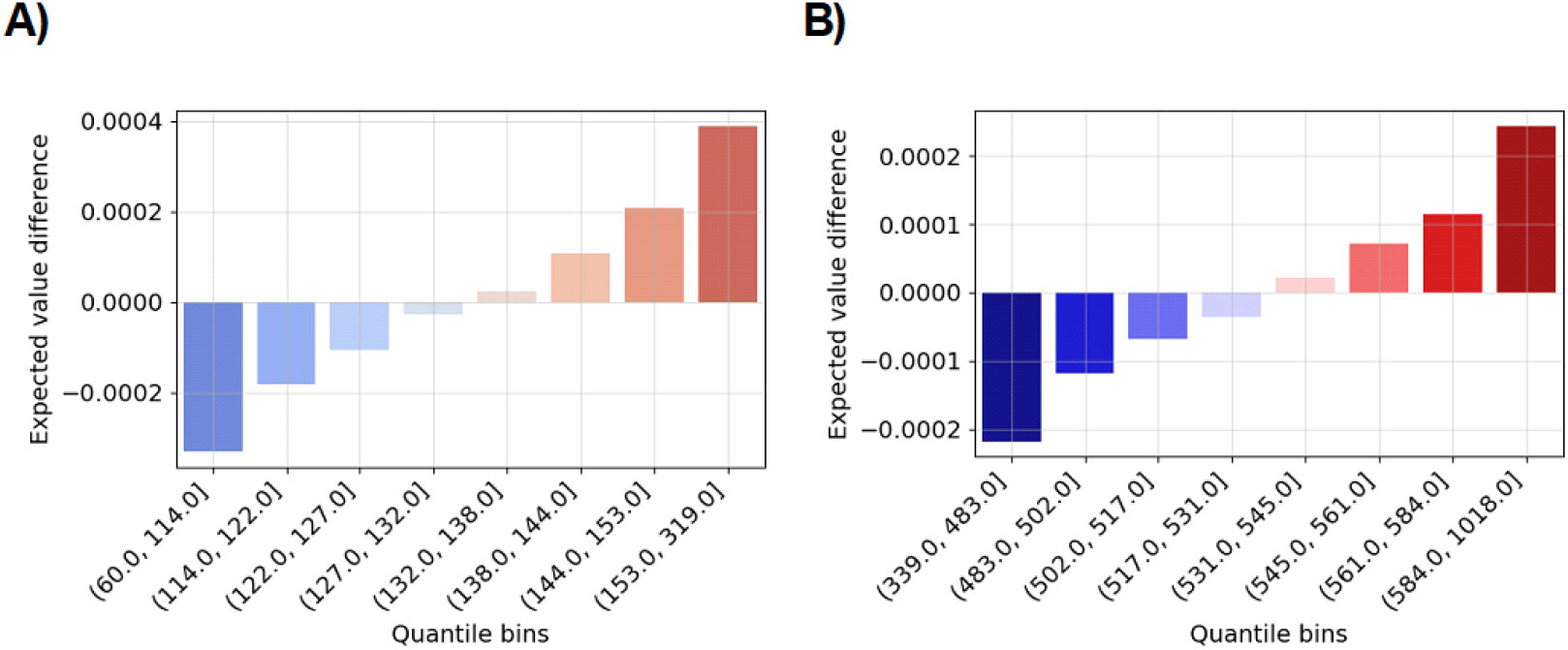
Monte Carlo verification of the conditional expectation condition in our central limit theorem. Bar charts show the difference between the proportion of simulations where two specific haplotypes share an IBD segment longer than 0.03 Morgans and the true success probability (y-axis). This statistic is stratified into eight quantile bins based on the total number of long IBD segments (x¬-axis). Sample sizes are A) two hundred and B) four hundred diploid individuals. The population size is ten thousand diploid individuals. The expectation is 132.78 in A) and 531.78 in B).

**Table S1:** Summary statistics of IBD graphs for the three phases of exponential growth (G3) and the population bottleneck (BN) demographic scenarios. Network structures of interest are the number of edges (Edges), the degree of the largest components (Largest), the number of trees of order 2 and 3 (Tree-2 and Tree-3), and the number of complete components of degree 3 or more (Complete). Summary statistics are aggregated over at least six hundred thousand simulations. Shapiro-Wilk tests at the significance level 0.05 are performed with 1000 replicates for at least 600 simulations, and the proportion of rejected null hypotheses is reported as S.W.t. The sample size is five thousand diploid individuals. The Morgans length threshold is 0.03.

| **Type** | **Structure** | **Avg** | **Var** | **Min** | **Max** | **S.W.t.** |
| --- | --- | --- | --- | --- | --- | --- |
| G3 | Edges | 3,085.66 | 12,827.06 | 2,554.00 | 3,716.00 | 0.29 |
|  | Largest | 15.60 | 9.22 | 9.00 | 50.00 | 1.00 |
|  | Tree2 | 757.96 | 635.16 | 644.00 | 880.00 | 0.08 |
|  | Tree3 | 99.71 | 94.29 | 54.00 | 149.00 | 0.42 |
|  | Complete | 251.55 | 230.65 | 185.00 | 3,118.00 | 0.14 |
| BN | Edges | 587.73 | 694.55 | 469.00 | 716.00 | 0.12 |
|  | Largest | 4.32 | 0.39 | 3.00 | 11.00 | 1.00 |
|  | Tree2 | 473.24 | 393.50 | 377.00 | 574.00 | 0.09 |
|  | Tree3 | 9.66 | 9.57 | 0.00 | 27.00 | 1.00 |
|  | Complete | 35.53 | 34.76 | 11.00 | 488.00 | 1.00 |

**Table S2:** Summary statistics of IBD graphs for different selection coefficients and the three phases of exponential growth demographic scenario. There is directional selection with different selection coefficients *s* ∈ [0.01, 0.02, 0.03, 0.4]. The same description of IBD graph features as in Table 1. Shapiro-Wilk tests at the significance level 0.05 are performed with 250 replicates for 150 simulations, and the proportion of rejected null hypotheses reported as S.W.t. The sample size is five thousand diploid individuals. The Morgans length threshold is 0.03.

| Type | Structure | Avg | Var | Min | Max | S.W.t. |
| --- | --- | --- | --- | --- | --- | --- |
| $s = 0.01$ | Edges | 3,407.38 | 21,526.32 | 2,916.00 | 4,143.00 | 0.36 |
|  | Largest | **24.33** | 50.80 | 11.00 | 89.00 | 0.97 |
|  | Tree2 | 737.77 | 626.12 | 636.00 | 842.00 | 0.05 |
|  | Tree3 | 95.81 | 92.23 | 57.00 | 138.00 | 0.07 |
|  | Complete | 242.41 | 215.76 | 187.00 | 305.00 | 0.05 |
| $s = 0.02$ | Edges | 4,693.51 | 140,436.48 | 3,579.00 | 8,212.00 | 0.95 |
|  | Largest | **73.97** | 1,219.95 | 22.00 | 346.00 | 0.97 |
|  | Tree2 | 697.19 | 588.38 | 596.00 | 791.00 | 0.10 |
|  | Tree3 | 86.65 | 83.70 | 53.00 | 126.00 | 0.09 |
|  | Complete | 220.37 | 199.88 | 161.00 | 281.00 | 0.10 |
| $s = 0.03$ | Edges | 8,242.12 | 2,283,864.57 | 4,998.00 | 37,933.00 | 0.97 |
|  | Largest | **230.39** | 12,224.19 | 39.00 | 819.00 | 0.97 |
|  | Tree2 | 659.10 | 565.21 | 562.00 | 759.00 | 0.07 |
|  | Tree3 | 78.43 | 74.69 | 46.00 | 119.00 | 0.11 |
|  | Complete | 199.95 | 181.88 | 145.00 | 254.00 | 0.06 |
| $s = 0.04$ | Edges | 16,486.56 | 24,295,227.62 | 7,747.00 | 72,775.00 | 0.97 |
|  | Largest | **484.92** | 38,683.32 | 89.00 | 1,229.00 | 0.97 |
|  | Tree2 | 630.68 | 529.35 | 546.00 | 731.00 | 0.02 |
|  | Tree3 | 72.95 | 70.26 | 41.00 | 108.00 | 0.11 |
|  | Complete | 185.76 | 167.85 | 135.00 | 241.00 | 0.07 |

**Table S3:**
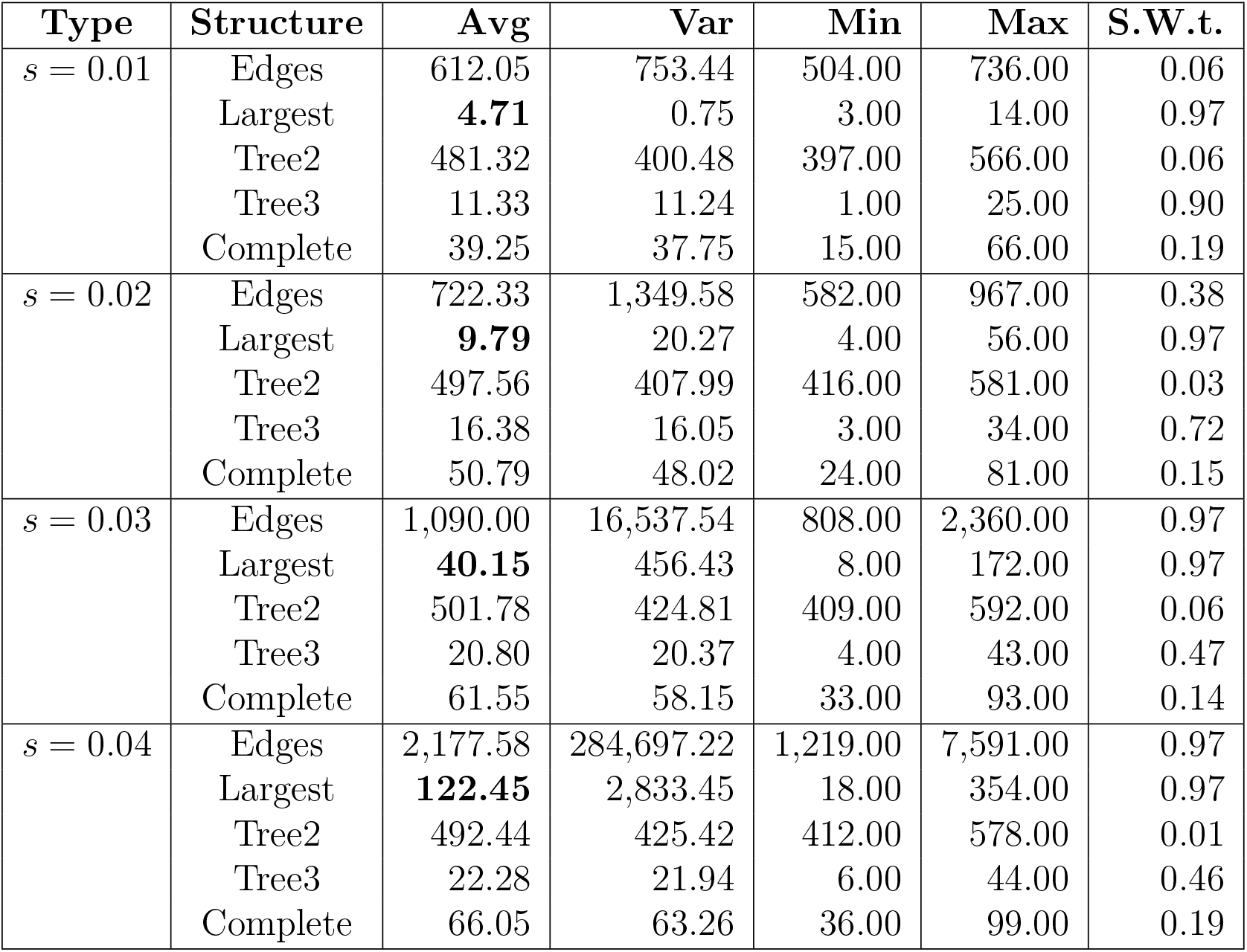
Summary statistics of IBD graphs for different selection coefficients and the population bottleneck demographic scenario. There is directional selection with different selection coefficients *s* ∈ [0.01, 0.02, 0.03, 0.4]. The same description of IBD graph features as in Table 1. Shapiro-Wilk tests at the significance level 0.05 are performed with 250 replicates for 150 simulations, and the proportion of rejected null hypotheses reported as S.W.t. The sample size is five thousand diploid individuals. The Morgans length threshold is 0.03.

| Type | Structure | Avg | Var | Min | Max | S.W.t. |
| --- | --- | --- | --- | --- | --- | --- |
| $s = 0.01$ | Edges | 612.05 | 753.44 | 504.00 | 736.00 | 0.06 |
|  | Largest | **4.71** | 0.75 | 3.00 | 14.00 | 0.97 |
|  | Tree2 | 481.32 | 400.48 | 397.00 | 566.00 | 0.06 |
|  | Tree3 | 11.33 | 11.24 | 1.00 | 25.00 | 0.90 |
|  | Complete | 39.25 | 37.75 | 15.00 | 66.00 | 0.19 |
| $s = 0.02$ | Edges | 722.33 | 1,349.58 | 582.00 | 967.00 | 0.38 |
|  | Largest | **9.79** | 20.27 | 4.00 | 56.00 | 0.97 |
|  | Tree2 | 497.56 | 407.99 | 416.00 | 581.00 | 0.03 |
|  | Tree3 | 16.38 | 16.05 | 3.00 | 34.00 | 0.72 |
|  | Complete | 50.79 | 48.02 | 24.00 | 81.00 | 0.15 |
| $s = 0.03$ | Edges | 1,090.00 | 16,537.54 | 808.00 | 2,360.00 | 0.97 |
|  | Largest | **40.15** | 456.43 | 8.00 | 172.00 | 0.97 |
|  | Tree2 | 501.78 | 424.81 | 409.00 | 592.00 | 0.06 |
|  | Tree3 | 20.80 | 20.37 | 4.00 | 43.00 | 0.47 |
|  | Complete | 61.55 | 58.15 | 33.00 | 93.00 | 0.14 |
| $s = 0.04$ | Edges | 2,177.58 | 284,697.22 | 1,219.00 | 7,591.00 | 0.97 |
|  | Largest | **122.45** | 2,833.45 | 18.00 | 354.00 | 0.97 |
|  | Tree2 | 492.44 | 425.42 | 412.00 | 578.00 | 0.01 |
|  | Tree3 | 22.28 | 21.94 | 6.00 | 44.00 | 0.46 |
|  | Complete | 66.05 | 63.26 | 36.00 | 99.00 | 0.19 |

## Footnotes

1 Functions that satisfy the Delta method conditions [10]: the first derivative at the expected number of detectable IBD segment around a locus exists and is nonzero.

2 The expected number of edges should be the same, if not for some approximations [36, 49].

3 The DRC and the detectable IBD rate have both been used in selection scans [5, 50]. Intuitively, they capture a similar population genetics signal.

